# Global reference mapping and dynamics of human transcription factor footprints

**DOI:** 10.1101/2020.01.31.927798

**Authors:** Jeff Vierstra, John Lazar, Richard Sandstrom, Jessica Halow, Kristen Lee, Daniel Bates, Morgan Diegel, Douglas Dunn, Fidencio Neri, Eric Haugen, Eric Rynes, Alex Reynolds, Jemma Nelson, Audra Johnson, Mark Frerker, Michael Buckley, Rajinder Kaul, Wouter Meuleman, John A. Stamatoyannopoulos

## Abstract

Combinatorial binding of transcription factors to regulatory DNA underpins gene regulation in all organisms. Genetic variation in regulatory regions has been connected with diseases and diverse phenotypic traits^1^, yet it remains challenging to distinguish variants that impact regulatory function^2^. Genomic DNase I footprinting enables quantitative, nucleotide-resolution delineation of sites of transcription factor occupancy within native chromatin^3–5^. However, to date only a small fraction of such sites have been precisely resolved on the human genome sequence^5^. To enable comprehensive mapping of transcription factor footprints, we produced high-density DNase I cleavage maps from 243 human cell and tissue types and states and integrated these data to delineate at nucleotide resolution ~4.5 million compact genomic elements encoding transcription factor occupancy. We map the fine-scale structure of ~1.6 million DHS and show that the overwhelming majority is populated by well-spaced sites of single transcription factor:DNA interaction. Cell context-dependent cis-regulation is chiefly executed by wholesale actuation of accessibility at regulatory DNA versus by differential transcription factor occupancy within accessible elements. We show further that the well-described enrichment of disease- and phenotypic trait-associated genetic variants in regulatory regions^1,6^ is almost entirely attributable to variants localizing within footprints, and that functional variants impacting transcription factor occupancy are nearly evenly partitioned between loss- and gain-of-function alleles. Unexpectedly, we find that the global density of human genetic variation is markedly increased within transcription factor footprints, revealing an unappreciated driver of cis-regulatory evolution. Our results provide a new framework for both global and nucleotide-precision analyses of gene regulatory mechanisms and functional genetic variation.

## Introduction

Genome-encoded recognition sites for sequence-specific DNA binding proteins are the atomic units of eukaryotic gene regulation. Binding of regulatory factors to their cognate elements in vivo shields them from nuclease attack, giving rise to protected single nucleotide-resolution DNA ‘footprints’. The advent of DNA footprinting using the non-specific nuclease DNase I^7^ marked a major turning point in analysis of gene regulation, and facilitated the identification of the first mammalian sequence-specific DNA binding proteins^8^. Genomic DNase I footprinting^3^ enables genome-wide detection of DNA footprints chiefly within regulatory DNA regions, but also over other genomic elements where DNase I cleavage is sufficiently dense.

DNase I footprints define sites of direct regulatory factor occupancy on DNA and can be used to discriminate sites of direct vs. indirect occupancy within orthogonal data from chromatin immunoprecipitation and sequencing (ChIP-seq) experiments. Cognate transcription factors (TFs) can be reliably assigned to DNase I footprints based on matching to consensus sequences, enabling TF-focused analysis of gene regulation and regulatory networks^9^, and the evolution of regulatory factor binding patterns within regulatory DNA^10^. DNase I is a small enzyme, roughly the size of a typical transcription factor that recognizes the minor groove of DNA and hydrolyzes single-stranded cleavages that, in principle, reflect both the topology and the kinetics of DNA-protein interaction. Previous efforts to exploit this feature^4^ were complicated by slight sequence-driven variation in cleavage preferences; however, these have now been exhaustively determined^11^, setting the stage for fully resolved tracing of DNA-protein interactions within regulatory DNA.

Currently we lack a comprehensive, nucleotide-resolution annotation of small DNA elements encoding regulatory factor recognition sites that are selectively occupied in different cell types. Such a reference is essential both for analysis of cell-selective occupancy patterns, and for systematic integration with genetic variation, particularly that associated with diseases and phenotypic traits. Here we combine sampling of >67 billion DNase I cleavages from >240 human cell types and states to index, with unprecedented accuracy and resolution, human genomic footprints and thereby the sequence elements that encode transcription factor recognition sites. We leverage this index to comprehensively assign footprints to transcription factor archetypes, define patterns of cell-selective occupancy, and analyze the distribution and impact of human genetic variation on regulatory factor occupancy and the genetics of common diseases and traits.

### Comprehensive mapping of human TF footprints

To create comprehensive maps of TF occupancy, we selected and deeply-sequenced high-quality DNase I libraries from 243 cell and tissues types derived from diverse primary cells and tissues (*n*=151), primary cells in culture (*n*=22), immortalized cell lines (*n*=10) and cancer cell lines and primary samples (*n*=60) (**Extended Data Table 1**). Collectively, we uniquely mapped 67.6 billion DNase I cleavage events (mean 278.2 million cleavages mapped per biosample), greatly eclipsing prior studies^4^. On average, 49.7% DNase I cleavage within each biosample mapped to DNase I hypersensitive sites covering 1-3% of the genome.

To identify DNase I footprints genome-wide, we developed a novel computational approach incorporating both chromatin architecture and exhaustively determined empirical DNase I sequence preferences to determine expected per-nucleotide cleavage rates across the genome, and to derive, for each biosample, a statistical model for testing whether observed cleavage rates at individual nucleotides deviated significantly from expectation (**Extended Data Fig. 1**, **Extend Data Fig. 2**, and **Methods**). We note that deriving cleavage variability models for each biosample accounts for additional sources of technical variability beyond DNase I cleavage preference.

Using this model, we performed *de novo* footprint discovery independently on each of the 243 biosamples, detecting on average 657,029 high-confidence footprints per biosample (range 220,580-1,664,065) (empirical false discovery rate <1%; **Methods**), and collectively 159.6 million footprint events across all biosamples. At the level of individual nucleotides, *de novo* footprints were highly concordant between replicates of the same cultured cell type or between same primary cell and tissues types from different individuals (median Pearson’s *r* = 0.83 and 0.74, respectively) (**Extended Data Fig. 3a-c)**. The significance of protected nucleotides tracked closely both the presence of known transcription factor recognition sequences and the level of per-nucleotide evolutionary conservation (**Extended Data Fig. 3d-e**). Within each biosample, genomic footprints encompassed an average of ~7.6 Mb (0.2%) of the genome, with a mean of 4.3 footprints per DHS with sufficient read depth for robust detection (normalized cleavage density within DHS ≥1).

### A unified index of human genomic footprints

Comparative footprinting across many cell types has the potential to illuminate both the structure and function of regulatory DNA, yet a systematic approach for joint analysis of genomic footprinting data has been lacking. Given the scale and diversity of the cell types and tissues surveyed, we sought to develop a framework that could integrate hundreds of available genomic footprinting datasets to increase the precision and resolution of footprint detection and, furthermore, serve as a scaffold to build a common reference index of TF-contacted DNA genome-wide.

To accomplish this, we implemented an empirical Bayes framework that estimates the posterior probability that a given nucleotide is footprinted by incorporating a prior on the presence of a footprint (determined by footprints independently identified within individual datasets) and a likelihood model of cleavage rates for both occupied and unoccupied sites (**Fig. 1a** and **Methods**). **Fig. 1b** depicts per-nucleotide footprint posterior probabilities computed for two DHS within the *RELB* locus across all 243 biosamples exposing the nucleotide-resolved TF occupancy architecture for each element. A notable feature of these data is the remarkable positional stability and discrete nature of footprints within each DHS across the tens to hundreds of biosamples. Indeed, plotting individual nucleotides scaled by their footprint prevalence across all samples precisely demarcates the core recognition sequences of diverse TFs (**Fig. 1b**, bottom).

**Figure 1:**
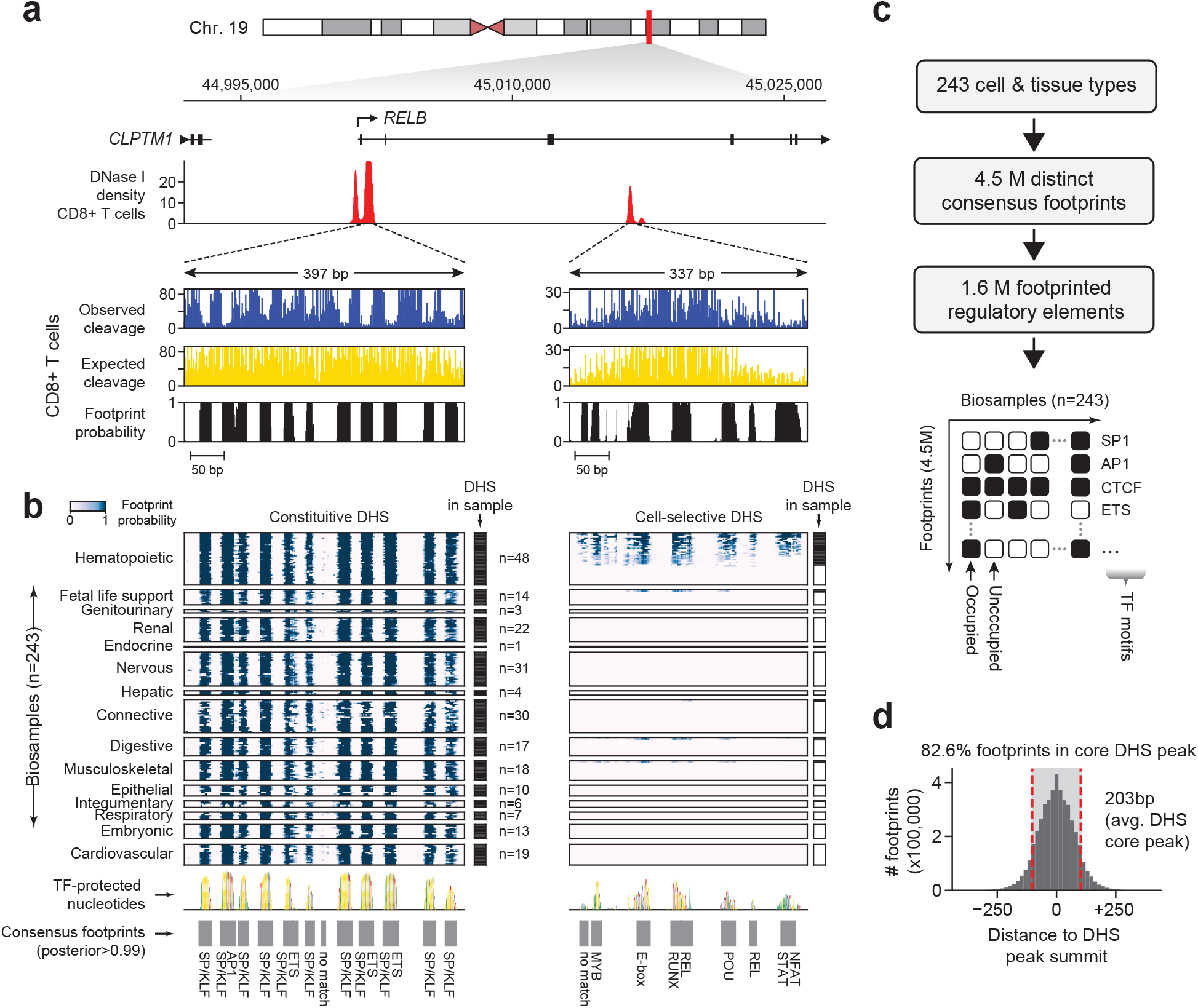
A nucleotide resolution atlas of transcription factor occupancy within the human genome. **a,** DNase I cleavage at regulatory DNA elements within the *RELB* locus in CD8+ T cells. Top, windowed DNase I cleavage density. Below, per-nucleotide cleavage counts and genomic footprint posterior probabilities at two DNase I hypersensitive sites in CD8+ T cells. **b**, Heatmap of genomic footprint posterior probabilities computed by integrating 243 datasets within the DHSs. Rows and columns correspond to individual biosamples and nucleotides grouped tissue/organ systems. Black ticks left of heatmap indicate whether region is DHS in each sample. Below, genome sequence scaled by footprint prevalence. Grey boxes define consensus footprints present in one or more cell and/or tissue types (footprint posterior>0.99). **c**, A consensus map of TF occupancy was derived from 243 cell and tissues covering 1.6 million DHS results in a comprehensive annotation of cis-regulatory DNA. **d**, Histogram of footprint location relative to DHS peak summit. Dashed red lines represent the average size of a DHS peak (203 bp).

To systematically create a reference set of TF-occupied DNA elements genome-wide, we applied the Bayesian approach to all DHSs detected within one or more of the 243 biosamples, and applied the same consensus approach used to establish the consensus DHS index^12^ to collate overlapping footprinted regions across individual biosamples into distinct high-resolution consensus footprints (**Methods**). Collectively, we delineated ~4.46 million consensus footprints within ~1.6 million individual DHS (**Fig. 1c**). 82.6% of consensus footprints localize directly within the core of a DHS peak (avg. width 203bp) with virtually all residing within 250bp of a DHS peak summit (**Figure 1d**). As expected, consensus-defined footprints were markedly more reproducible than footprints detected independently within a given biosample (avg. Jaccard distance 0.43 vs 0.29, respectively) (**Extended Data Fig. 4d-e**). Consensus footprints had an average width of 16 bp (middle 95%: 7–44 bp; 90%: 7-36 bp; 50%: 9-21 bp), and collectively annotate 2.1% (72 Mb) of the human genome reference sequence, a compartment slightly larger than protein coding sequences (~1.5%).

### Assigning footprints to TF motifs

Our understanding of the recognition motif landscape of human transcription factors has undergone dramatic development during the past decade, and recognition sequences now exist for all major families and subfamilies, and for a large number of individual TF isoforms^13–16^. We thus sought to create a reference mapping between annotated transcription factors and consensus human genomic footprints by (i) compiling and clustering all publicly available motif models^13,17,18^; (ii) creating non-redundant TF archetypes by placing closely-related TF family members on a common sequence axis (**Extended Data Fig. 5**, **Extended Data Table 2** and **Methods**); (iii) aligning these archetypes to the human reference sequence at high stringency (*p*<10^−4^); and (iv) enumerating all potential TF archetypes compatible with each consensus footprint on the basis of overlap and match stringency (**Methods**). In total, 80.7% of the ~4.46 million consensus footprints could be assigned to at least one TF recognition sequence (≥90% overlap; **Methods**), of which 860,780 (19.3%) could be unambiguously assigned to a single factor, and 2,038,220 (45.7%) to a single factor with two lower-ranked alternatives (**Extended Data File 2**).

### Primary architecture of regulatory regions

Despite intensive efforts over the past three decades the primary architecture of regulatory regions has remained elusive, with the singular exception of the interferon ‘enhanceosome’^19^. A prerequisite for understanding the primary architecture of active regulatory DNA is accurate tracing of the TF:DNA interface over an extended interval. Because transcription factor engagement within DNA major or minor grooves creates subtle alterations in DNA shape and protects underlying phosphate bonds from nuclease attack via steric hindrance^5^, we asked to what extent fluctuations in corrected DNase I cleavage rates within consensus footprints accurately reflect the topology of the TF:DNA interface. Poly-zinc fingers are the most prevalent class of human transcription factors and have recognition interfaces that potentially cover tens of nucleotides^16^. The DNA recognition domain of the genomic master regulator CTCF comprises 11 zinc fingers, potentially encoding 33bp of sequence (or DNA shape^20^) recognition. We identified 25,852 footprints that coincided precisely with CTCF motifs. Transposing the average corrected per-nucleotide cleavage propensity with an extended co-crystal structure of CTCF^21^ accurately traced all features of the protein:DNA interaction interface, including focal hypersensitivity within bent hinge region between zinc fingers 7 and 9^22,23^ (**Fig. 2a** and **Methods**). A similar result was obtained for widely divergent classes of DNA binding domains such as the paired-box containing TF PAX6^24^ (**Fig. 2b**) and other TFs with extant co-crystal structures (not shown). Critically, these topological features are evident at the level of individual TF footprints on the genome (**Fig. 2a-b** and **Extended Data Fig. 6**), indicating that the extended profile of corrected per-nucleotide DNase I cleavage across entire regulatory regions should, in principle, provide a snapshot of the primary structure of active regulatory DNA.

**Figure 2:**
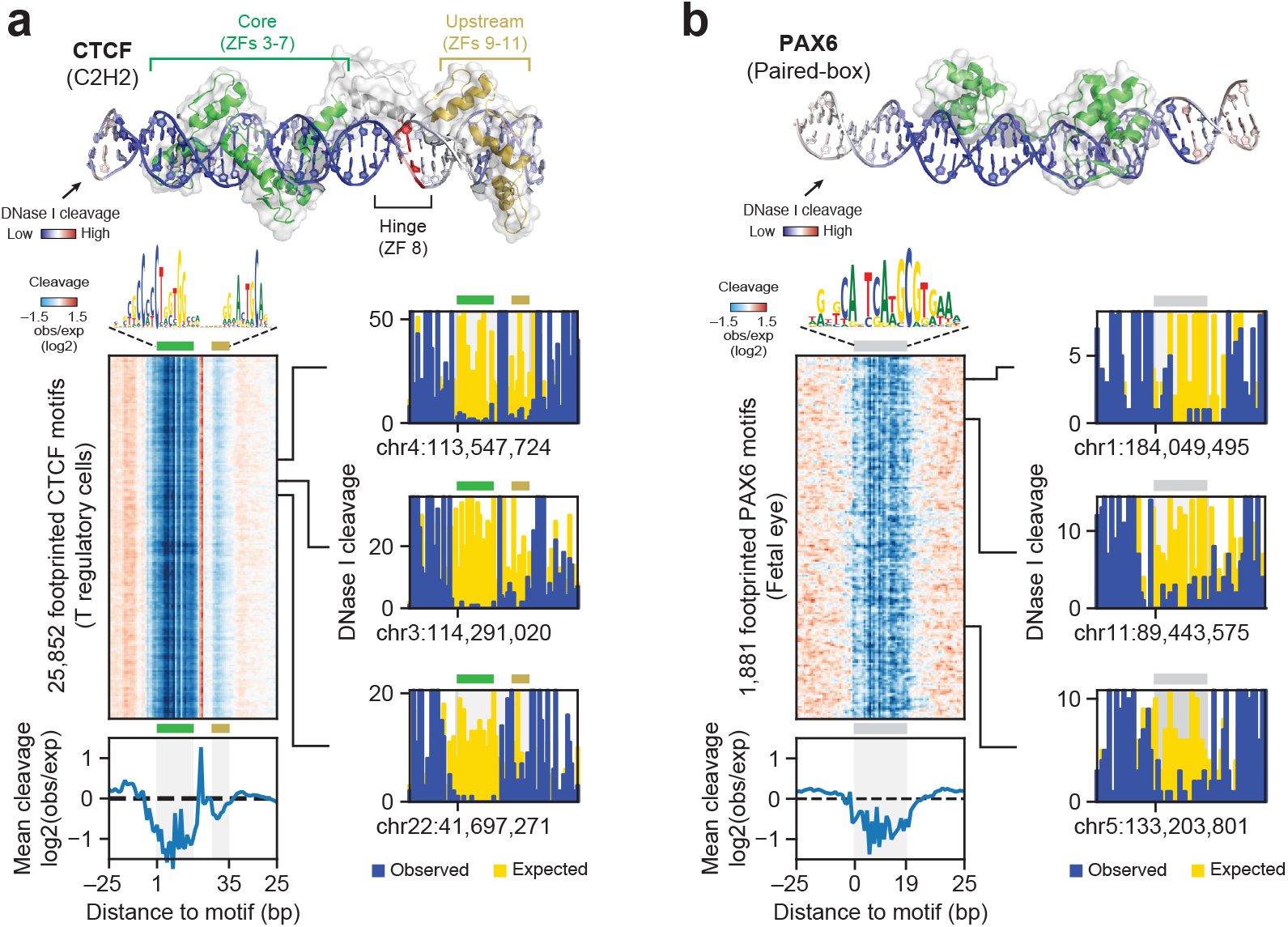
DNase I footprints reflect the topological structure of individual TF:DNA interactions. **a**, Top, Physical structure of CTCF zinc fingers 3-11 bound to its cognate DNA recognition sequence (PDB: 5YEF and 5YEL)^21^. DNA is colored by mean ratio of observed vs. expected cleavages at footprinted CTCF motifs in T regulatory cells. Left, heatmap of a relative cleavage at each of the 25,852 footprinted CTCF motifs. Bottom, aggregate DNase I cleavage summed over all CTCF footprints. Right, DNase I cleavage (observed and expected) at three randomly selected footprints. **b**, Same as **a** for paired-box transcription factor PAX6 (PDB: 6PAX)^24^.

### Distinguishing independent vs. cooperative modes of TF occupancy

Transcription factors compete cooperatively with nucleosomes for access to regulatory DNA^25,26^. Although fundamental to eukaryotic gene regulation, it is currently unknown whether nucleosome-enforced TF cooperativity derives primarily from local protein-protein interactions or results from the synergistic effect of independent TF:DNA binding events^26^. We reasoned that the number, relative spacing, and morphology of TF binding events within individual regulatory elements could be used to gain insight into the mechanistic basis of TF cooperativity. We observed that the average footprint width for diverse TFs tightly corresponded to the total width of its recognition sequence (Spearman’s ρ=0.82, *p*-value=0.001) indicating that DNase I cleavage precisely delineates the boundaries between occupied and unoccupied DNA at nucleotide resolution (**Fig. 3a**).

**Figure 3:**
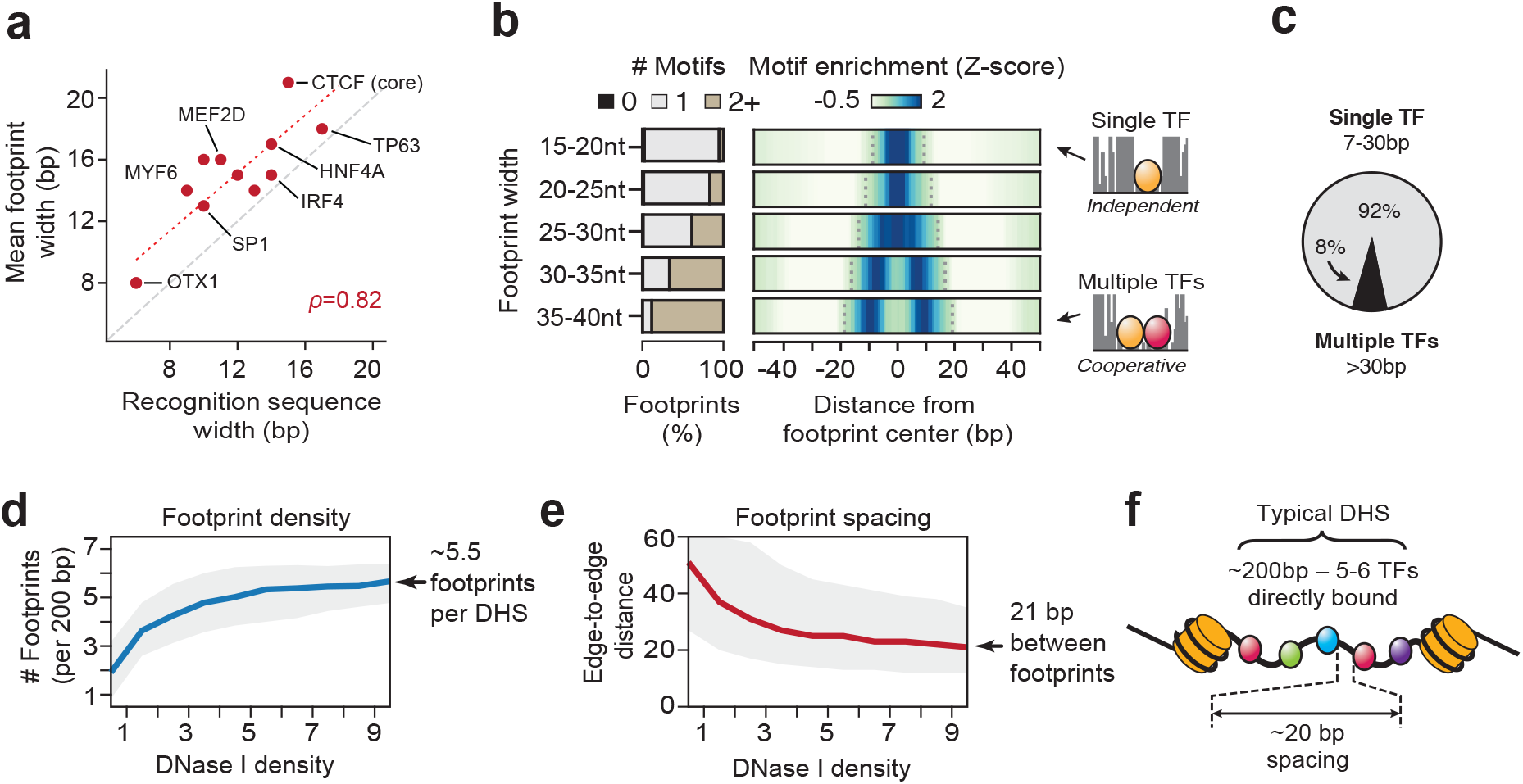
Distinguishing modes of transcription factor occupancy within regulatory DNA. **a,** The width of footprints for diverse TFs tightly correlates with the width of their recognition sequence (Spearman’s *ρ*=0.82, *p*-value=0.001). **b**, Overlap and spatial enrichment of TF recognition sequences within footprints binned by width. Left, Proportion of footprints uniquely overlapped by 0, 1 or 2 or more recognition sequences. Right, density heatmap of motif occurrences around footprints binned by width. **c,** Percentage of footprints that likely represent the occupancy of a single TF (≤30bp) or multiple TFs (>30bp). **d–e,** Footprint density and footprint spacing (distance edge-to-edge) vs. average DNase I density within DHSs. Grey indicates the middle 50%-ile. **f,** A typical regulatory element (DHS) harbors ~5-6 directly bound TFs spaced roughly 20-bp from each other.

Since the width of genomic footprints tightly tracks the physical structure of individual TFs bound to DNA (**Fig. 2a-b**), and direct TF:TF interactions are dependent on close proximity^19^, as such interactions should be reflected in larger footprints that harbor multiple TF recognition sites. Conversely, independent TF:DNA interaction events should be reflected by compact and widely-spaced footprints harboring single TF recognition sites. As such, the prevalence of cooperativity mediated by direct TF:TF interactions vs. synergy of independent binding events should be reflected in relative proportion of wide multi-motif footprints vs. well-spaced single footprints. Larger footprints are overwhelmingly associated with two (or more) recognition sequences (**Fig. 3b**), yet such footprints represent only 8% of the global footprint landscape. By contrast, 92% of footprints harbor a single TF recognition site (**Fig. 3c**).

Transcription factors can distort DNA upon engagement; as such, the spacing of transcription factors can be critical for establishing an active regulatory structure. To quantify global footprint spacing patterns, we first binned each DHS by its average accessibility across all samples (because footprint discovery depends on total DNase I cleavage; **Extended Data Fig. 1b**), and for each bin we computed the mean number of footprints present per element and their relative edge-to-edge spacing. The density of footprints within the most deeply sampled DHSs genome-wide plateaued at an average of 5.5 per 200 bp (average width of a DHS peak) (**Fig. 3d**), which is in remarkable agreement with a theoretical prediction of the number of human TFs required to destabilize a canonical nucleosome^26^ and to encode specificity^27^. Within DHSs, footprints exhibit an average edge-to-edge spacing of ~21bp (middle 50%, 12-35bp) (**Fig. 3e**). Taken together, these results are compatible with the observed lack of evolutionary constraint on the spacing and orientation of TF motifs^28^ and provide strong evidence that regulatory DNA marked by DHSs is chiefly instantiated by independent but synergistic TF binding modes (**Fig. 3f**).

### Cell-selective TF occupancy patterns

Analysis of footprint occupancy across all biosamples revealed strong enrichment for the recognition sequences of key regulatory TFs in their cognate lineages (**Extended Fig. 7a** and **Methods**; for example: HNF4A in fetal intestine and liver; GATA factors in erythroid and placental/trophoblast cells and tissues; NEUROG1 in brain; myogenic regulatory factors (e.g, MYF6, MYOD, etc.) in muscle and lung; MEIS1 in developing eye, brain, and muscle tissues; and PAX6 in fetal eye). In total, we identified 609 motif models matching footprinted sequences (**Methods**); these models encompassed 64 distinct archetypal transcription factor recognition codes (**Extended Data Table 2**), representing virtually all major DNA-binding domain families. For degenerate motifs where the same sequence is recognized by many distinct TFs, we observed highly cell-selective occupancy patterns that could be decomposed into coherent groups corresponding with cell type and function (**Extended Data Fig. 7b-d**).

### Most DHSs encode a single regulatory topology

Given that a significant fraction of DHSs are shared across two or more cell/tissue types^12,29^, we next asked whether differential TF occupancy within the same regulatory DNA region (vs. differential actuation of entire DHSs) could be a major driver of cell-selective regulation. Nucleotide-resolution DNase I cleavage provides a topological fingerprint of each DHS, reflecting its unique combination and ordering of occupying TFs. Although detectable on manual inspection^4^, systematic analysis of differential TF occupancy has previously not been possible due to dominance of intrinsic cleavage propensities when many data sets are combined. To enable unbiased detection of differential footprint occupancy, we developed a statistical framework to test for differences in relative DNase I cleavage rates at individual nucleotides across many samples, analogous with methods developed for the identification of differentially expressed genes (**Methods**). To estimate the proportion of differentially regulated footprints within DHSs of a given cell/tissue type, we compared footprint occupancy within DHSs broadly accessible in both nervous-system derived samples (*n*=31) with non-nervous-system derived samples (*n*=212). We selected 67,368 DHSs that were highly accessible in at least 10 nervous and non-nervous derived samples, and for each DHS, performed a per-nucleotide differential test (**Fig. 4a,b** and **Extended Data Fig. 8**). This analysis identified only a small proportion of DHSs (1,720 DHSs; 2.5%) containing a differentially footprinted element (**Fig. 4c**). Most of these DHSs harbored a single differentially regulated footprint, while a small fraction contained 2-4 differentially occupied elements (**Fig. 4c**). Nonetheless, differentially occupied footprints were significantly enriched recognition sites for known nervous system regulators such as REST, NFIB, ZIC1, and EBF1 (**Fig. 4c**, bottom right and **Extended Data Fig. 9**) and tissue-selective occupancy events paralleled expression of nearby genes (in the case of REST occupancy) (**Extended Data Fig. 10**).

**Figure 4:**
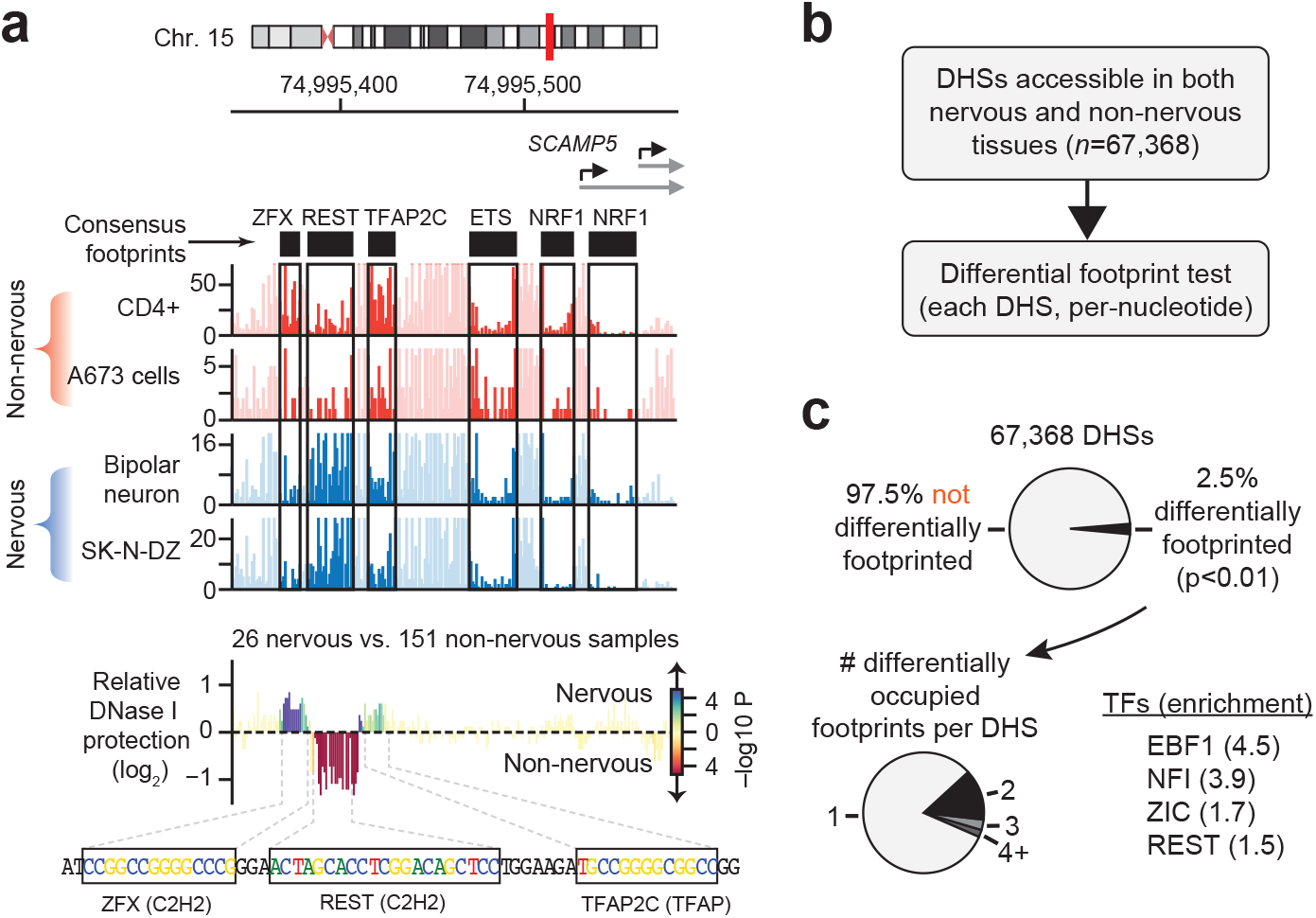
Comparative footprinting identifies cell-selective TF occupancy at nucleotide resolution. **a,** Comparative footprinting within the *SCAMP5* promoter identifies 3 footprints differentially occupied in nervous cell and tissue types. Top, DNase I cleavage in two exemplar nervous and non-nervous cell types. Bottom, mean differential per nucleotide cleavage(log2) between nervous-system derived (*n*=26) and non-nervous samples (*n*=12). The color of each bar indicates the statistical significance (–log10 *p*) of the per-nucleotide differential test. **b,** Differential footprint testing within thousands of DHS accessible in between nervous and non-nervous related biosamples. **c**, The vast majority of tested DHSs encode a single TF binding topology. Top, percentage of the DHSs tested that containing one or more differentially occupied element. Bottom left, distribution of differentially footprinted elements per DHS. Bottom right, selected TF recognition sequences significantly enriched in differentially occupied footprints (binomial test p<0.01). Indicated in parenthesis is the fold-enrichment vs. expected (based on prevalence of footprinted motif in tested regions).

Collectively, the above results indicate that the vast majority of regulatory DNA regions marked by DHSs encode a single structural topology reflecting a fixed pattern of footprint occupancy. Nonetheless, at a small minority of elements, DHSs provide a scaffold for cell context-specific TF occupancy that is typically confined to a single or small number of footprinted elements.

### Functional variants in TF footprints

Identifying genetic variants likely to impact regulatory function has remained challenging^2^. Deep sequence coverage at DHSs enables *de novo* genotyping of regulatory variants and simultaneous characterization of their functional impact on local chromatin architecture by quantifying and comparing DNase I cleavage for each allele of a given element^2,4^. The 243 biosamples we analyzed were derived from 147 individuals. *De novo* genotyping of all samples (**Methods**) revealed 3.76 million single nucleotide variants within DHSs, of which 1,656,597 were heterozygous and had sufficient read depth (≥35 overlapping reads) to accurately quantify allelic imbalance.

Across individuals, we conservatively identified 117,626 chromatin altering variants (CAVs) that impacted DNA accessibility on individual alleles (median 2.4-fold imbalance) (**Fig. 5a**, **Extended Data Fig. 11a** and **Methods**). Within DHSs, CAVs were markedly enriched in core consensus footprints, even after controlling for the increased detection power within this compartment (**Fig. 5b** and **Extended Data Fig. 12**).

**Figure 5:**
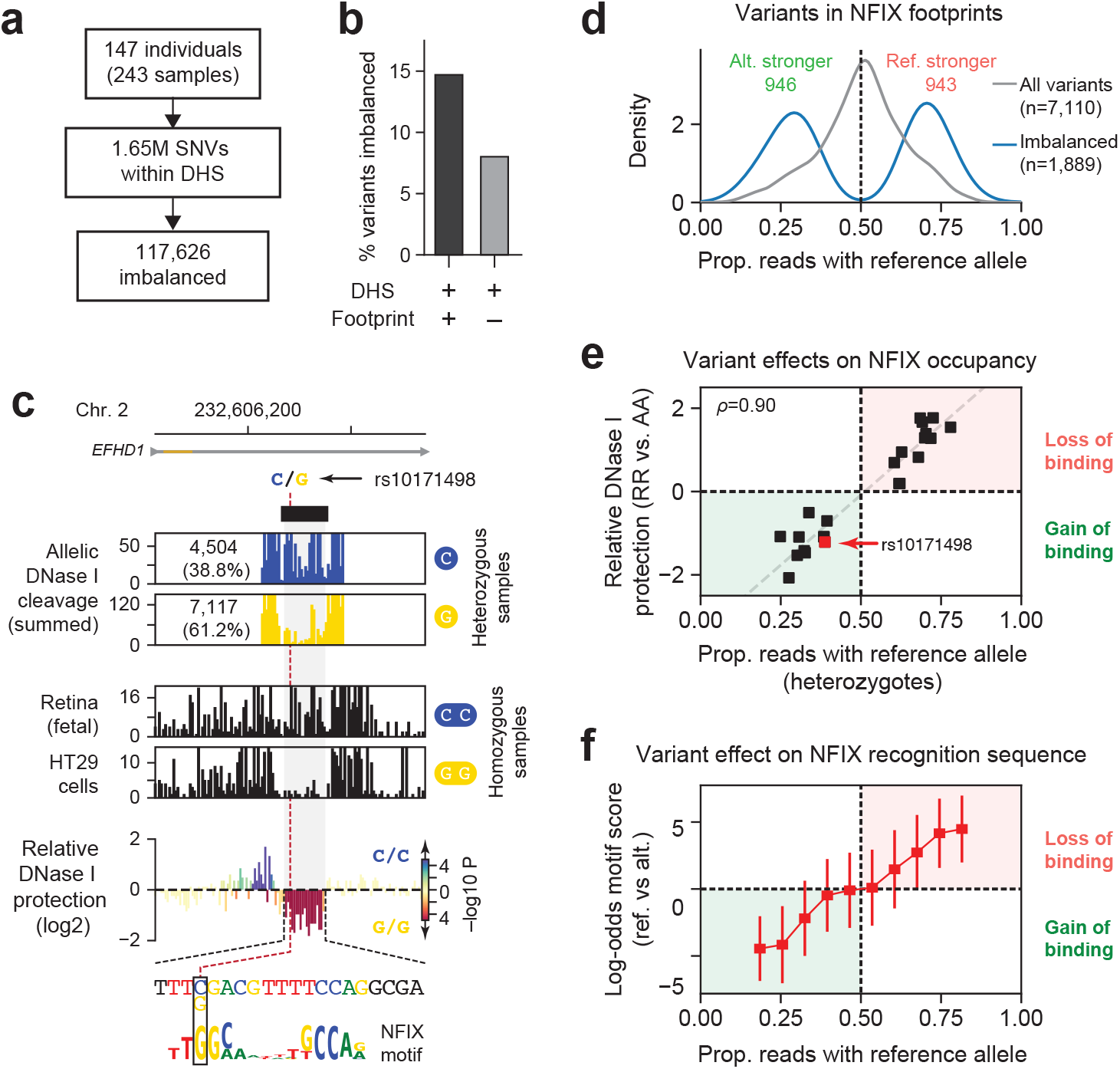
Functional genetic variation localizes to TF footprints. **a**, Allelic imbalance was assessed at all variants overlapping a DNase I footprints (consensus footprint probability < 0.01). **b**, Percentage of variants imbalanced in DNase I footprints and DHS peaks (but not in footprints). **b**, Variant rs10171498 results in the gain of a NFIX footprint. Top, allelically resolved per-nucleotide DNase I cleavage aggregated from 56 heterozygous samples. Middle, DNase I cleavage in two selected samples homozygous for either reference or alternative alleles. Bottom, mean differential per nucleotide cleavage (log2) between homozygous reference (*n*=74) and alternative samples (*n*=12). The color of each bar indicates the statistical significance (–log10 *p*) of the per-nucleotide differential test (**Methods**). The variant and differentially footprinted nucleotides precisely colocalize to a NFIX recognition element. **d**, Density histogram of allelic ratios for variants overlapping a footprinted NFIX recognition sequence. Grey, all variants tested for imbalance (*n*=7,110). Blue, all variants significantly (n=1,889) imbalanced. **g**, Scatterplot of allelic imbalance computed from heterozygous individuals (*x*-axis) vs. the relative difference in footprint depth between homozygous individuals at variants overlapping an NFIX footprint. Each point pertains to a SNV within a footprinted NFIX binding site imbalanced in heterozygotes and differentially footprinted in homozygotes. Grey line indicates fit linear model. **e**, Allelic imbalance measurements parallels predicted energetic effects of variants within NFIX footprints. Shown is the mean log-odds motif score (reference vs. alternate allele) of all tested variants within footprinted motifs binned by allelic ratios.

In protein-coding regions, most functional genetic variation is expected to be deleterious, with rare gain-of-function alleles^30^. Protein-DNA recognition interfaces are likewise presumed to be susceptible to disruption at critical nucleotides, predisposing to loss-of-function alleles. Strikingly, we found CAVs to be nearly evenly partitioned between loss-(disruption of binding) and gain-of-function (increased or *de novo* binding) alleles (**Fig. 5c-d** and **Extended Data Fig. 10c**). Homozygosity for either the reference or alternative allele paralleled results from heterozygotes and further revealed that structural changes due to TF occupancy were precisely confined to the DNA sequence recognition interface (**Fig. 5c**, bottom). In many cases, SNVs detected in both heterozygous and homozygous configurations showed strong agreement between allelic ratios and relative footprint strength (**Fig. 5e**; Spearman’s *ρ*=0.9, *p*-value < 10^−5^). Variants residing within core recognition motifs in footprints were markedly enriched for imbalance vs. non-footprinted motifs; were localized to high-information content positions within the recognition interface (**Fig. 5c**, bottom and **Extended Data Fig. 13**); and paralleled the predicted energetic effect of the variant on the TF binding site (**Fig. 5f** and **Extended Data Fig. 14**), thus providing a direct quantitative readout of functional variation impacting TF occupancy.

### DNA elements encoding footprints are hypermutable

We next explored the global distribution of human genetic variation relative to consensus footprints. Transcription factor binding sites appear to be gradually remodeled over evolutionary time via sequential small mutations^31^ that could ultimately affect function and phenotype^32^. However, patterns of genetic variation within regulatory DNA have not been characterized with high precision. To quantify these, we calculated nucleotide diversity (π) within and around consensus genomic footprints using whole-genome sequencing data compiled from >65,000 individual under the TOPMED project^33^ (**Methods**). Canonically, reduced levels of π reflect the elimination of deleterious alleles from the population by natural selection, and hence are typically indicative of functional constraint^34^. Surprisingly, we found a dramatic increase in nucleotide diversity centered precisely within the core of genomic footprints (**Fig. 5a**), and thus that these elements are highly polymorphic in human populations. This result eclipses prior global analyses indicating that transcription factor occupancy sites are generally not under substantial purifying selection^4^ both in the magnitude of the observed effect, and in its nucleotide-precise localization within the core of genomic footprints.

The focal increase in genetic diversity within footprints indicated that the nucleotides encoding footprinted elements may have an increased mutational load vs. immediately adjacent sequences. To explore this possibility, we focused on variants with extremely low allele frequencies in human populations (minor allele frequency < 10^−4^); such variants are assumed to result from *de novo* germline (ie., non-segregating) mutation and are often used as a surrogate for mutation rate in humans^35^. We found that the distribution of extremely rare variants around and within genomic footprints mirrored that of nucleotide diversity, compatible with context-driven increased mutation rate in the sequences underlying footprints (**Fig. 5b**). Of note, many transcription factors favor recognition of dinucleotide combinations such as CpGs that are intrinsically hypermutable. Conversely, *de novo* mutations have been implicated in the genesis of TF recognition sites^36,37^. Thus, hypermutation within genomic footprints may fill a key evolutionary role by favoring variability in TF occupancy and hence natural variation in gene regulation.

### GWAS variants are enriched in TF footprints

Given the above, genetic variation within genomic footprints should, in principle, be a key contributor phenotypic variation; however, to date this has defied accurate quantification. We therefore next resolved the large set of variants strongly associated (nominal *p*-value < 5×10^−8^) with diverse diseases and phenotypic traits from the NHGRI/EBI GWAS Catalogue^38^ to consensus genomic footprints. To account for the baseline increase in genetic variation present within genomic footprints described above, we performed exhaustive (1,000x) sampling matched variants (by minor allele frequency, linkage-disequilibrium (LD) structure, and distance to the nearest gene) from the 1,000 Genome Project^39^ (**Methods**). Additionally, we expanded both GWAS catalogue and matched sampled variants to include variants in perfect LD (*r*^2^=1). Within DHSs, aggregated GWAS catalogue SNPs were enriched within footprints but not in non-footprinted subregions, and enrichment within footprints increased monotonically with footprint strength (**Fig. 6c** and **Extended Data Fig. 15**).

**Figure 6:**
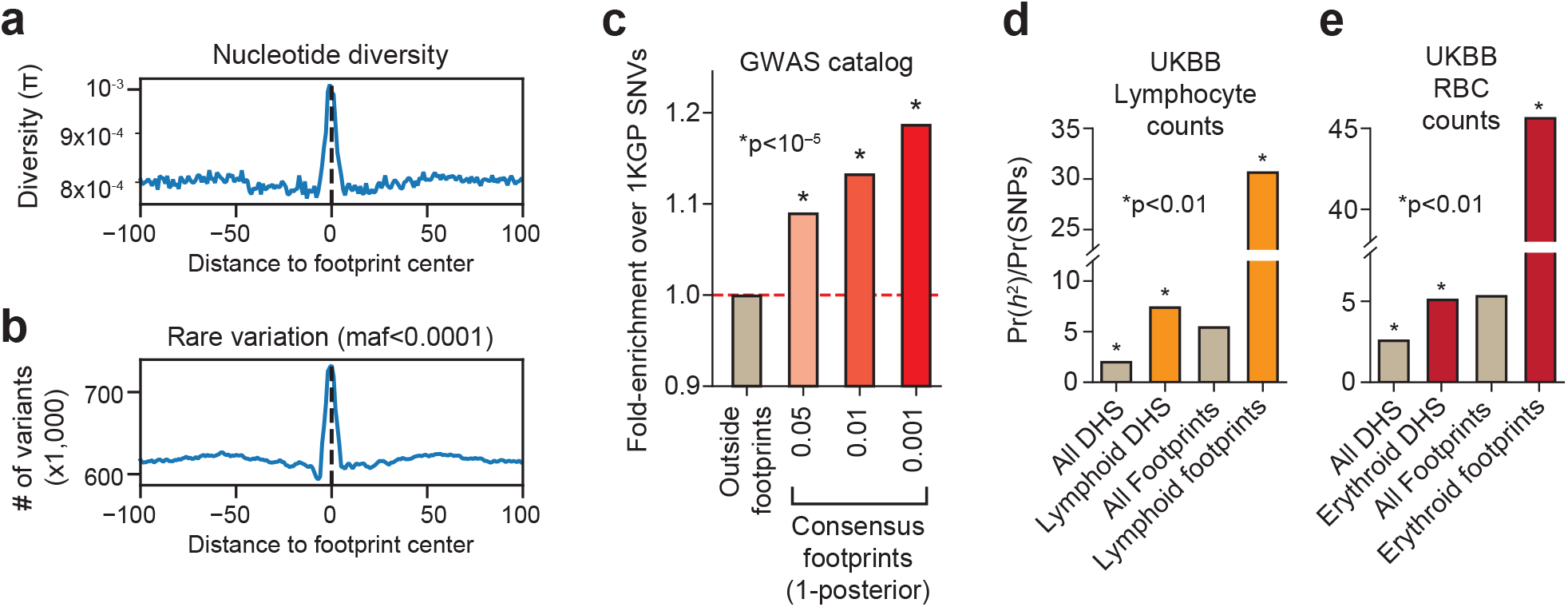
Human genetic variation is broadly enriched within genomic footprints. **a,** Distribution of genetic variation with respect to consensus footprints. Plotted is the mean per nucleotide diversity determined from whole genome sequencing of 62,784 individuals (TOPMED project). **b,** Histogram of the distribution of rare variation (minor allele frequency<0.0001) within and surrounding genomic footprints. **c,** Enrichment of GWAS variation within or outside consensus genomic footprints over randomly sampled variants from 1,000 Genome Project. Enrichment was computed after expanding both GWAS and sampled variants with those in perfect LD (*r*^2^=1.0, central European population). **d,** Enrichment of SNP-based trait heritability using LD-score regression for UK BioBank GWAS traits lymphocyte counts (**c**) and red blood cell counts (**d**). Asterisk denotes statistically significant enrichments (* indicates p-value<0.01).

The GWAS catalogue aggregates hundreds of traits, with corresponding expected diversity in cognate cell/tissue types. To gain a more accurate view of the enrichment of trait-associated variants in footprints, we compared SNP-based trait heritability of individual traits^40,41^. Using summary statistic data from individual GWAS studies from the UK BioBank, we applied partitioned LD-score regression to compute the relative heritability contribution of variants within all DHSs and footprints collectively vs. DHSs and footprints therein from the expected cognate cell type for the trait (**Fig. 6d-e**). This analysis revealed striking enrichment of variants that account for trait heritability in footprints generally (>5-fold) and most prominently in footprints from the cognate cell type (up to >45-fold) (**Fig. 6d-e**). Collectively, we conclude that the genetic signals from disease- and trait-associated variants within DHSs emanate from transcription factor footprints, and that variants within footprints are major contributors to trait heritability.

## Discussion

Our report details the highest resolution view to date of regulatory factor occupancy patterns on the human genome measured across an expansive range of cell and tissue contexts sampled from >140 genotype backgrounds. The scale and breadth of the data have enabled the delineation of a reference set of ~4.5 million genomic sequence elements that collectively define nucleotides critical for genome regulation and function and form the building blocks of regulatory DNA. These footprints now provide a ready and extensible nucleotide-precise reference for diverse analyses, particularly those involving genetic variation.

The preferential localization of disease- and trait-associated variation within regulatory DNA has heretofore been described in terms of entire regulatory regions demarcated by DHSs or clusters thereof. Our results now show that genetic association and heritability signals from regulatory DNA overwhelmingly emanate from indexed transcription factor footprints, which should greatly facilitate the connection of disease- and trait-associated genetic variation with genome function.

Perhaps most strikingly, we report that human genetic variation is itself concentrated within transcription factor footprints, owing apparently to a combination of mutation propensity and the evolved sequence recognition repertoire of human transcription factors, which favors hypermutable nucleotide combinations (e.g., CG dinucleotides). Given that human and mouse TFs share the large majority of their recognition landscapes, concentration of variation within TF occupancy sites has likely played a considerable role in shaping human – and indeed all mammalian – gene regulation. It implies, furthermore, that the genome is heavily primed for regulatory evolution, providing a possible mechanism underlying facilitated phenotypic evolution^42^

## Author contributions

J.V. designed experiments and performed analysis. J.L. aided in the design of statistical methods for footprint detection. R.S. organized and aided in the processing of raw data. W.M. aided in the creation of the consensus footprint index. J.S. and J.V. wrote the paper. All others participated in data generation and organization.

## Acknowledgements

This work was supported by NHGRI grant U54HG007010 to J.A.S

## References

1. Maurano, M. T. et al. Systematic localization of common disease-associated variation in regulatory DNA. Science 337, 1190–1195 (2012).

2. Maurano, M. T. et al. Large-scale identification of sequence variants influencing human transcription factor occupancy in vivo. Nat. Genet. 47, 1393–1401 (2015).

3. Hesselberth, J. R. et al. Global mapping of protein-DNA interactions in vivo by digital genomic footprinting. Nat. Methods 6, 283–289 (2009).

4. Neph, S. et al. An expansive human regulatory lexicon encoded in transcription factor footprints. Nature 489, 83–90 (2012).

5. Vierstra, J. & Stamatoyannopoulos, J. A. Genomic footprinting. Nat. Methods 13, 213–221 (2016).

6. Roadmap Epigenomics Consortium et al. Integrative analysis of 111 reference human epigenomes. Nature 518, 317–330 (2015).

7. Galas, D. J. & Schmitz, A. DNAse footprinting: a simple method for the detection of protein-DNA binding specificity. Nucleic Acids Res. 5, 3157–3170 (1978).

8. Dynan, W. S. & Tjian, R. The promoter-specific transcription factor Sp1 binds to upstream sequences in the SV40 early promoter. Cell vol. 35 79–87 (1983).

9. Neph, S. et al. Circuitry and dynamics of human transcription factor regulatory networks. Cell 150, 1274–1286 (2012).

10. Stergachis, A. B. et al. Conservation of trans-acting circuitry during mammalian regulatory evolution. Nature 515, 365–370 (2014).

11. Lazarovici, A. et al. Probing DNA shape and methylation state on a genomic scale with DNase I. Proc. Natl. Acad. Sci. U. S. A. 110, 6376–6381 (2013).

12. Meuleman, W., Muratov, A., Rynes, E., Halow, J. & Lee, K. Index and biological spectrum of accessible DNA elements in the human genome. BioRxiv (2019).

13. Jolma, A. et al. DNA-binding specificities of human transcription factors. Cell 152, 327–339 (2013).

14. Yin, Y. et al. Impact of cytosine methylation on DNA binding specificities of human transcription factors. Science 356, (2017).

15. Najafabadi, H. S. et al. C2H2 zinc finger proteins greatly expand the human regulatory lexicon. Nat. Biotechnol. 33, 555–562 (2015).

16. Lambert, S. A. et al. The Human Transcription Factors. Cell 175, 598–599 (2018).

17. Khan, A. et al. JASPAR 2018: update of the open-access database of transcription factor binding profiles and its web framework. Nucleic Acids Res. 46, D1284 (2018).

18. Kulakovskiy, I. V. et al. HOCOMOCO: towards a complete collection of transcription factor binding models for human and mouse via large-scale ChIP-Seq analysis. Nucleic Acids Res. 46, D252–D259 (2018).

19. Panne, D., Maniatis, T. & Harrison, S. C. An atomic model of the interferon-beta enhanceosome. Cell 129, 1111–1123 (2007).

20. Rohs, R. et al. The role of DNA shape in protein-DNA recognition. Nature 461, 1248–1253 (2009).

21. Yin, M. et al. Molecular mechanism of directional CTCF recognition of a diverse range of genomic sites. Cell Res. 27, 1365–1377 (2017).

22. Arnold, R., Burcin, M., Kaiser, B., Muller, M. & Renkawitz, R. DNA bending by the silencer protein NeP1 is modulated by TR and RXR. Nucleic Acids Res. 24, 2640–2647 (1996).

23. MacPherson, M. J. & Sadowski, P. D. The CTCF insulator protein forms an unusual DNA structure. BMC Mol. Biol. 11, 101 (2010).

24. Xu, H. E. et al. Crystal structure of the human Pax6 paired domain-DNA complex reveals specific roles for the linker region and carboxy-terminal subdomain in DNA binding. Genes & Development vol. 13 1263–1275 (1999).

25. Svaren, J., Klebanow, E., Sealy, L. & Chalkley, R. Analysis of the competition between nucleosome formation and transcription factor binding. J. Biol. Chem. 269, 9335–9344 (1994).

26. Mirny, L. A. Nucleosome-mediated cooperativity between transcription factors. Proc. Natl. Acad. Sci. U. S. A. 107, 22534–22539 (2010).

27. Wunderlich, Z. & Mirny, L. A. Different gene regulation strategies revealed by analysis of binding motifs. Trends Genet. 25, 434–440 (2009).

28. Vierstra, J. et al. Mouse regulatory DNA landscapes reveal global principles of cis-regulatory evolution. Science 346, 1007–1012 (2014).

29. Thurman, R. E. et al. The accessible chromatin landscape of the human genome. Nature 489, 75–82 (2012).

30. Fu, W. et al. Analysis of 6,515 exomes reveals the recent origin of most human protein-coding variants. Nature 493, 216–220 (2013).

31. Payne, J. L. & Wagner, A. The robustness and evolvability of transcription factor binding sites. Science 343, 875–877 (2014).

32. Prud’homme, B. et al. Repeated morphological evolution through cis-regulatory changes in a pleiotropic gene. Nature 440, 1050–1053 (2006).

33. Taliun, D., Harris, D. N., Kessler, M. D. & Carlson, J. Sequencing of 53,831 diverse genomes from the NHLBI TOPMed Program. BioRxiv (2019).

34. Kellis, M. et al. Defining functional DNA elements in the human genome. Proc. Natl. Acad. Sci. U. S. A. 111, 6131–6138 (2014).

35. Carlson, J. et al. Extremely rare variants reveal patterns of germline mutation rate heterogeneity in humans. Nat. Commun. 9, 3753 (2018).

36. He, X. et al. Methylated Cytosines Mutate to Transcription Factor Binding Sites that Drive Tetrapod Evolution. Genome Biol. Evol. 7, 3155–3169 (2015).

37. Zemojtel, T., Kielbasa, S. M., Arndt, P. F., Chung, H.-R. & Vingron, M. Methylation and deamination of CpGs generate p53-binding sites on a genomic scale. Trends Genet. 25, 63–66 (2009).

38. Buniello, A. et al. The NHGRI-EBI GWAS Catalog of published genome-wide association studies, targeted arrays and summary statistics 2019. Nucleic Acids Res. 47, D1005–D1012 (2019).

39. 1000 Genomes Project Consortium et al. A map of human genome variation from population-scale sequencing. Nature 467, 1061–1073 (2010).

40. Finucane, H. K. et al. Partitioning heritability by functional annotation using genome-wide association summary statistics. Nat. Genet. 47, 1228–1235 (2015).

41. Bulik-Sullivan, B. K. et al. LD Score regression distinguishes confounding from polygenicity in genome-wide association studies. Nat. Genet. 47, 291–295 (2015).

42. Gerhart, J. & Kirschner, M. The theory of facilitated variation. Proc. Natl. Acad. Sci. U. S. A. 104 Suppl 1, 8582–8589 (2007).

