## Extended Figures, Data and Methods for "Global reference mapping and dynamics of human transcription factor footprints"

### Extended Data for "Global reference mapping and dynamics of human transcription factor footprints"


**Methods** 3

Digital genomic footprinting data 3

DNase I sequence preference model 3

Computing expected per-nucleotide cleavage rates 3

Derivation of cleavage dispersion model 4

Per-nucleotide cleavage testing 4

Bayesian modeling 5

Derivation of consensus footprints 6

Testing for differential DNase I cleavage 6

Evolutionary conservation 6

Transcription factor binding site predictions 6

Clustering TF recognition sequences by similarity 7

Assignment of TF recognition sequences to consensus footprints 7

Co-crystal structural modeling 7

Human genetic variation 7

Genotyping pipeline 8

Detecting allelic imbalance 8

Enrichment of imbalanced variants within TF recognition sequences 9

Energetic effects of variation on TF recognition sequences 9

GWAS variation 9

Stratified LD-score regression analysis 9

**Data availability** 10

**Code availability** 10

**Extended Data Figures** 11

Extended Data Figure 1. Statistical modeling of DNase I cleavage variation 11

Extended Data Figure 2. Footprint detection within in single dataset 11

Extended Data Figure 3. Genomic footprints are reproducible. Harbor evolutionary constrained nucleotides, and overlap TF recognition sequences 11

Extended Data Figure 4. Empirical Bayes framework identifies footprints with markedly increase reproducibility 12

Extend Data Figure 5. Clustering motifs by similarity 12

Extended Data Figure 6. Aggregate DNase I cleavage profiles for diverse transcription factors. 12

Extended Data Figure 7. Cell-selective occupancy of TF recognition sequences 12

Extended Data Figure 8. Differential occupancy within the UCP2 promoter in human brain and neuronal cells 12

Extend Data Figure 9. Differentially occupied nucleotides reflect aggregate DNase I cleavage profiles 13

Extended Data Figure 10. Expression of genes nearby differentially occupied REST sites is enriched in brain and neuronal cell types 13
Extended Data Figure 11. Detection and quantification of allelic imbalance 13
Extended Data Figure 12. Allelically imbalanced variants are enriched within consensus footprints. 13
Extended Data Figure 13. Enrichment of imbalanced variants within footprinted TF recognition sequences. 14
Extended Data Figure 14. Allelic imbalance parallels the predicted energetic effect of the variant on the TF binding site 14

**Extended Data Tables 14**
Extended Data Table 1. Overview and summary of DGF data used in this study 14
Extended Data Table 2. Clustering of TF recognition sequence models 14

**Extended Data Files 14**
Extended Data File 1. Footprints identified in individual datasets 14
Extended Data File 2. Consensus-defined footprints across 243 biosamples 15
Extended Data File 3. Posterior probability matrix of consensus footprints by biosamples 15

**Extended References 16**

### Methods

##### Digital genomic footprinting data

The digital genomic footprinting datasets used in this study were released as part of the ENCODE[1] and Roadmap Epigenomics Consortia[2]. DNase I digestion, purification of small double-hit fragments and sequencing library preparation was performed as in ref. 3. Sequencing was performed on the HiSeq platform (Illumina). Raw sequencing reads were trimmed to remove adapter sequences and aligned to the human genome (hg38/GRCh38) using bwa[4] version 0.7.12) with the following parameters: "`-Y -l 32 -n 0.04`" and "`-n 10 -a 750`" for alignment and mate-pairing (aln and sampe, respectively). DHS peaks (i.e., hotspots) were determined using (hotspot2; http://github.com/Altius/hotspot2).

##### DNase I sequence preference model

A sequence model of DNase I preference was constructed as in refs. 5,6. Briefly, deproteinated genomic DNA derived from IMR90 (fetal lung fibroblasts) cells was digested with bovine DNase I (Sigma-Aldrich). DNase I-released fragments isolated, processed and sequenced as described above. For each uniquely mapping sequencing tag, the 5' alignment position, $i$, was used to extract the DNA sequence hexamer covering positions $[i-3, i+2]$ with respect to the strand of the alignment. The total amount of each hexamer was normalized by the total number of hexamers (considering both strands) within the 36-bp uniquely mappable genome.

##### Computing expected per-nucleotide cleavage rates

Expected cleavage rates are computed by redistributing cleavage counts to the DNase I sequence preference model described above. First, the total cleavages (*w*) within a ±5 bp of each base (11 bp window) genome-wide is calculated such that:

$$w_i = \sum_{j=i-5}^{i+5} n_j$$

From these values a sliding trimmed mean is computed (removing the top and bottom 1%) in ±50 bp intervals (101 bp window) which represents an estimate of total expected cleavages within ±5 bp of each nucleotide.

$$w_i^s = \overline{(w_{i-50}, \dots, w_{i+50})}_{0.01}$$

Next, the underlying hexamer sequence for each nucleotide, *i*, is used to create any array of relative sequence preference across the genome (a). These relative preference values are then normalized in ±5 bp sliding windows (as above).

$$p_i = \frac{a_i}{\sum_{j=i-5}^{i+5} a_j}$$

Finally, the expected cleavages for each nucleotide ($n_i'$) is determined by multiplying the expected cleavages in a ±5 bp window to its normalized relative preference value at the same position.

$$n_i' = p_i w_i^s$$

An expected count is derived independently for each strand and are combined to generate a final aggregated expected count.

##### Derivation of cleavage dispersion model

Statistical detection of footprints uses a properly fit per-nucleotide cleavage dispersion model from which to test whether observed cleavage rates significantly deviate from the expected. A dispersion model is created independently from the observed data in three steps:

1. **Compute expected cleavage rates.** Expected cleavage rates are generated from the observed cleavages rates (as above), except without windowed smoothing (i.e., $n_i' = p_i w_i$)

2. **Fit parameters for statistical distribution.** For each predicted cleavage rate all of the observed rates at their corresponding sites are collected and the parameters of a negative binomial distribution are fitted to the observed rates by maximum likelihood estimation

3. **Local smoothing of model parameters.** As the predicted cleavage rate increases the number of observations decreases (due to a preponderance of low cleavage rates). To reduce the noise in fitting negative binomial distributions to each predicted cleavage rate and to estimate the NB parameters in the case of insufficient data, a piecewise linear regression is performed to determine the parameters with respect to the predicted cleavage rate. The resulting regression coefficients are used to compute the negative binomial parameters in all subsequent analysis.

##### Per-nucleotide cleavage testing

The significance of cleavage rate deviation at a nucleotide *i* is determined by comparing its observed cleavages to the cleavage rate expected at an unoccupied nucleotide. First, per-nucleotide deviation *p*-values are computed from the lower-tail of a negative binomial distribution parameterized by μ and *r* from the dispersion model with respect to the expected cleavage rate. The lower tail of the negative binomial is computed using the incomplete beta function. A footprint is defined by short stretches of nucleotides which display relative protection from cleavage due to protein engagement. In the null case (no footprint), the cleavage rates at adjacent nucleotides are independent (**Extended Data Fig. 2b**). As such, we test for joint local deviation of cleavage rates by combining adjacent per-nucleotide *p*-values in a local window (±3 bp; 7 bp total) using Stouffer's *Z*-score method.

Calibration of *p*-values to account for multiple testing is performed empirically by sampling cleavage counts from the observed variance in the dispersion model. Specifically, each the cleavage count for each nucleotide is resampled from the expected distribution per the dispersion model. Sampled nucleotides are then processed identical to the observed cleavage rates, such that per-nucleotide and windowed *p*-values are computed. This sampling procedure was performed 1,000 times independently and the *p*-values were aggregated to generate a reference null distribution. The observed *p*-values are sorted and ranked against the null distribution to yield an empirical false discovery rate.

##### Bayesian modeling

We formulated an Empirical Bayes framework that computes posterior *p*-values after considering all observed data. This method is used to generate a list of high-confidence reference footprints across hundreds of samples. The posterior probability (*p*) of a footprint at individual nucleotide is:

$$p(\theta_+|X) = \frac{P(\theta_+)\mathcal{L}(\theta_+|X)}{P(\theta_+)\mathcal{L}(\theta_+|X) + \big(1 - P(\theta_+)\big)\mathcal{L}(\theta_-|X)}$$

The footprint prior, $P(\theta_+)$, is the number of datasets that a nucleotide is found within a footprint (FDR<0.05) divided by the number datasets in which that nucleotide lays within a DNase I hypersensitive site (as defined by hotspot2). The likelihood function corresponding an unoccupied nucleotide, $\mathcal{L}(X|\theta_-)$, is the product of individual negative binomial probabilities corresponding to dispersion model parameterized by the expected cleavage rate ($\mu$, $r$).

$$\mathcal{L}(\theta_-|x_i) = \prod_{j=i-3}^{i+3} \binom{x_j + r_j - 1}{x_j} \big(1 - p_j\big)^{r_j} p_j^{x_j}$$

$$p_j = \frac{r_j}{r_j + \mu_j}$$

The likelihood function corresponding an occupied nucleotide, $\mathcal{L}(X|\theta_+)$, is generated by computing an expected rate of cleavage reduction at footprinted elements genome wide. First, for each dataset we fit a Beta distribution to the ratio of observed over expected cleavages (depletion ratio) at all FDR 5% footprints identified within individual datasets (capping the ratio values at 1.0). Then, for each nucleotide we re-estimate the depletion ratio by updating the Beta distribution (α'= α + observed, β'=(expected–observed) + α). These updated parameters are used to generate maximum *a posteriori* (MAP) estimates of the depletion ratio ($\mu_{MAP}$) and expected variation of this ratio ($\sigma^2_{MAP}$) at each nucleotide. A per-nucleotide footprint depletion estimate is then finally calculated from the average of the MAP mean estimates weighted by the inverse of the MAP standard deviation considering all datasets with an identified footprint at that nucleotide. The footprint depletion estimate is multiplied by the expected cleavage rate for unoccupied nucleotides to compute an expected cleavage count at an occupied base. The parameters for the negative binomial are then set per this value and the likelihood is computed.

##### Derivation of consensus footprints

We derived an index of footprints in the human genome by considering the total collection of all per-dataset footprints called at FDR level 1%. Briefly, for each genomic locus, we aligned the location and dispersion of footprints across datasets to delineate consensus coordinates supported by at least 50% of all footprint-contributing datasets. This approach is identical to the one used to delineate consensus DNase I hypersensitive sites (DHSs) as used in the accompanying manuscript by Meuleman *et al.*[7]

##### Testing for differential DNase I cleavage

We created a test whether modeling distributions of cleavage rates as two groups is more likely than modelling a data as a single group. The test models the $\log_2$ transformed observed vs. expected ratios as normal distributions with known mean but with uncertain variance estimates. As such, we model the cleavage rate variation using a Bayesian approach. We estimate prior hyperparameters on the nucleotide cleavage ratio variance using a scaled-inverse $\chi 2$ distribution from all positions within a DHS. The scaled-inverse $\chi 2$ distribution is a conjugate prior to the normal distribution in which the Student's t results as the predictive posterior distribution of the observed vs. expected cleavage rates. The likelihood of the data is the combination of the posterior probability distribution of the cleavage ratio with the likelihood the observed cleavage data:

$$p(X|u, v', \sigma^{2\prime}) = \sum_{i=1}^{n} \int \mathcal{L}(\theta_{-}|x_i)\, t_{v\prime}(\theta|\mu, \sigma^{2\prime}) d\theta$$

$$v' = v_o + n$$

$$\sigma^{2\prime} = \frac{v_o \sigma_0^2 + \sum_{i=1}^{n}(\phi_i - \mu)^2}{v_o + n}$$

where $v_o$ and $\sigma_0^2$ are the (hyper)parameters of the fitted scaled-inverse $\chi 2$ distribution, $\mu$ is the mean of the $\log_2$ transformed observed vs. expected cleavage ratios ($\phi$) and $n$ is the number of samples. The likelihood of the data is computed considering two groups or as a single group using numerical approximation. A log-likelihood ratio is computed as $LLR = L_{AB} - (L_A + L_B)$. A likelihood ratio test ($\chi^2$; 3 degrees of freedom) is used to test for statistical differences between the models (two groups vs. one group) and hence differential footprints. We combined adjacent per-nucleotide *p*-values in a local window (±3 bp; 7 bp total) using Stouffer's *Z*-score method.

##### Evolutionary conservation

The per-nucleotide phyloP[8] 100-way conservation track was downloaded from the UCSC Genome Browser corresponding to human genome build hg38/GRCh38 used for all analyses.

##### Transcription factor binding site predictions

TF recognition sequence models were obtained from a large scale SELEX[9], JASPAR (2018)[10], and HOCOMOCO[11] (version 11) and scanned genome-wide using the software package

MOODS[12] with the following parameters: "`--p-value 1e-4 --lo-bg 0.2977 0.2023 0.2023 0.2977`"

##### Clustering TF recognition sequences by similarity

To systematically collapse redundant motifs by similarity we used an approach similar to Maurano et al.[13]. Briefly, we used TOMTOM[14] to compute the distances between all motif pairs (2,179 motif models). We then performed hierarchical clustering using Pearson correlation as the distance metric and complete linkage. The tree was cut at height 0.7. For each of the 286 clusters, we then randomly selected a seed motif model to which we aligned all other motifs within cluster (both position and orientation).

##### Assignment of TF recognition sequences to consensus footprints

To assign TF recognition sequences to consensus footprints, we collapsed genome-wide transcription factor binding site predictions of all TF models by translating the coordinates and orientation of motif match into relative to its assigned cluster, removing redundant assignments of the same motif cluster at identical genomic coordinates. We then overlapped these collapsed motifs with consensus footprints, requiring that either 90% of motif is overlapped by footprint or vice-versa.

For all other analyses, all genome-wide TF recognition sequence matches were overlapped (≥3bp) with reference footprints (posterior probability>0.99).

##### Co-crystal structural modeling

To visualize DNase I cleavage with respect to the physical CTCF physical structure, we obtained protein-DNA co-crystals corresponding to the ZFs 2-8 and 6-11 (PDB 5YEF and 5YEL)[16] and assembled them together *in silico* to generate a full-length structural representation of the DNA-binding domain interface. For PAX6, we obtained the TF:DNA co-crystal (PDB:6PAX[25]) and computationally predicted and aligned the structure of an extended fragment of DNA using Web 3DNA 2.0 (http://web.x3dna.org)[26] (sequence: `GCTCCTCTTAAGCATTTTCACGCATGAGTGCACAGACCTTAAGA)`. Assembly, analysis and visualization of structural data was performed using PyMOL[17].

##### Human genetic variation

Estimates of human population genetic variation was obtained from NHLBI TOPMED project (freeze 5), downloaded from the Bravo webserver (http://bravo.sph.umich.edu). Only variants passing all filters ("PASS") were considered. Indels were removed from all analysis. Nucleotide diversity measurements (π) were calculated using whole genome sequences π for a heterozygous site is where $p_i$ is allele frequency:

$$\pi = 1 - \sum_{i=1}^{n} p_i^2$$

##### Genotyping pipeline

Genotype information was ascertained directly from the DNase I data using a standard genotyping pipeline. First genotypes from individual datasets were called directly from the BAM files generated for digitial genomic footprinting using bcftools (version 1.7) commands `mpileup` with the following parameters: "`-Q 20 -d 1000 -I -D -a FORMAT/DP,FORMAT/AD`" and `call` with the parameters " `-f GQ -cv -Ov`". Further filtering was performed using vcftools[24] (version 0.1.14) with settings "`--minQ 500 --minGQ 50 --minDP --max-alleles 2`". As multiple samples used as part of this project correspond to single individuals, we calculated relatedness (vcftools --relatedness) and merged all alignment files corresponding to single individuals to increase genotyping sensitivity. The merged data corresponding to 143 individuals (comprising 243 datasets) was genotyped as above. Finally, variants where filtered that significantly deviated from Hardy-Weinberg equilibrium (HWE exact test; $p<0.01$). After filtering 3,758,562 million SNVs heterozygous in one or more individuals remained.

##### Detecting allelic imbalance

For each sample, we filtered reads for which variants introduce mapping artifacts using the software package WASP[18]. Duplicate reads were discarded randomly. We determined allele-specific read counts for all heterozygous position further filtering for number of alignment mismatches (>1 reference; >2 alternate) and genotype quality (>20). In addition, reads for which the variant position within 3 bp of the 5' end was discarded to avoid any variants that may affect DNase I cleavage rates.

We used beta-binomial distribution to test for imbalance, which allows for additional parameter to model dispersion. To tune the parameters of distribution we selected a set high confidence SNVs (n=407,511) with strong statistical power (heterozygous in 2 or more samples, at least 20 reads covering either allele in each sample and at least 100 total reads summed over all samples) and for each SNV computed the mean and standard deviation of the allelic ratio across all samples. We set the parameters $\alpha$ and $\beta$ of the beta-binomial using the following equations:

$$\alpha = \mu \times \left(\frac{1}{\sigma^2} - 1\right)$$
$$\beta = (1-\mu) \times \left(\frac{1}{\sigma^2} - 1\right)$$

where $\mu$ and $\sigma$ correspond to the average of the means and standard deviations over all selected SNVs.

For each SNV containing ≥35 total summed reads over all heterozygous samples (from either allele) computed statistical significance of allelic imbalance using the Beta-binomial distribution (parameterized as above). Overall, 1,656,597 variants were tested for allelic imbalance. Multiple testing correction of the Beta-binomial *p*-values was performed using the Benjamini-Hochberg method. Due to the extremely conservative nature of our test, imbalanced variants were established at a false discovery rate of 20%.

##### Enrichment of imbalanced variants within TF recognition sequences

To assess the relative enrichment if imbalanced SNVs with respect to TF recognition sequences, we used an approach identical to ref. 13. Briefly, all tested variants were aligned relative to all motif models distinguishing whether each motif instance overlapped a consensus footprint (**Extended Data Table 2**). The proportion of SNVs imbalanced was computed using variants with an FDR<20% and reference allelic ratio ≥70%. Motifs were filtered to have ≥40 SNVs per position and at least 3 position with ≥7 imbalanced SNVs. Statistical significance of the enrichment was determined by permutation of the imbalance labels with respect to all SNVs within the motif and 20bp flanking regions. Multiple testing correction was performed using the Benjamini-Hochberg method and motifs were considered significant at a false discovery rate of 5%.

##### Energetic effects of variation on TF recognition sequences

To measure the effects of sequence variation within putative TF bindings sites with respect to allelic imbalance, we first created a custom genome containing all possible alleles by encoding variant positions using ambiguous IUPAC DNA codes (e.g., A/T = W). We then identified motif matches within the custom genome using the variant aware mode of MOODS[12] (see *Transcription factor binding site predictions*) that scans all possible alleles at ambiguous positions. For each variant that overlapped a motif instance, we computed a motif match score separately for each allele computed the log difference between the two scores.

##### GWAS variation

Disease- and trait-associated human genetic variation was obtained from the NHGRI-EBI GWAS Catalog[19] (v1.0 all associations) (https://www.ebi.ac.uk/gwas/docs/file-downloads). The enrichment of GWAS data within DHS or footprints was performed by an empirical sampling approach. Random SNVs from the 1,000 Genomes Project[20] (1KGP) (central European population) were sampled and overlapped with DHS or DNase I footprints. Sampled 1KGP SNVs were matched with observed GWAS SNVs for allele frequency and linkage disequilibrium structure (total number of SNVS with $r^2$=1 with GWAS allele). Both the GWAS and sampled SNPs were expanded to all SNVs in perfect LD before assessing overlaps with either DHS peaks or footprints. The sampling procedure was repeated 1,000 times to estimate the parameters of a normal distribution ($\mu$, $\sigma$). These parameters were used to calculate the upper-tail *p*-value of the observed overlap of GWAS SNPs.

##### Stratified LD-score regression analysis

SNP-based trait heritability was computed using LD-score regression (S-LDSC)[20,21] using a reference set of HapMap3 SNPs. In addition to the baseline set of 97 annotations provided as part of the LDSC software package (baseline-LD model v2.2; https://data.broadinstitute.org/alkesgroup/LDSCORE/), we created additional SNP annotations with respect to DHS and footprints. To generate these additional annotations, we merged all DHS or footprints with bedops[22] and lifted over their genomic coordinates to human genome build hg19 using CrossMap[23] with default parameters. GWAS summary statistics for two UK BioBank traits were download from (http://www.nealelab.is/uk-biobank; Benjamin Neale lab) corresponding to RBC count (30010_irnt.gwas.imputed_v3.both_sexes) and lymphocyte count (30120_irnt.gwas.imputed_v3.both_sexes). LDSC was run for each DHS or footprint annotation independently (each run including the 97 baseline annotations).

#### Data availability

All raw sequence data generated for this study can be accessed with GEO accession numbers found within **Extended Data Table 1**. DNase I hotspots and peaks are released as part of ENCODE Consortium and available for downloaded through the ENCODE data portal website (http://http://www.encodeproject.org/).

Footprint calls and other Extended Data Files are persistently hosted at ZENODO and can be publicly accessed at https://doi.org/10.5281/zenodo.3603548.

#### Code availability

Scripts and software used for analysis are available at http://www.github.com/jvierstra/footprint-tools

#### Extended Data Figures

##### Extended Data Figure 1. Statistical modeling of DNase I cleavage variation

**a,** A negative binomial model was fit from the distribution of observed cleavage counts for each predicted cleavage rate. Shown are histograms of observed cleavage counts in CD19+ B cells at all genomic sites with 5, 25, or 60 expected cleavages. Red, the negative binomial distribution fit to the observed data using maximum likelihood estimation (see **Methods**). Red curve, Poisson distribution with λ set to the corresponding expected cleavage rate. Lower right panel shows the means of fitted negative binomial distributions vs. means of observed cleavage rates. Dashed grey line indicates $y=x$ for reference. **b,** Estimated power of empirical cleavage dispersion model. Computed *p*-values for different cleavage rate effect sizes with respect to expected cleavage rates in CD19+ B cells. Colored lines represent the modeled effect size (depletion of cleavages) relative to the expected rate corresponding to a hexamer sequence model.

##### Extended Data Figure 2. Footprint detection within in single dataset

**a,** Example of footprint detection within promoters *TMEM143* and *SYNGR4* in CD19+ B cells. Expected cleavages are generated by reassigning observed cleavages according to a hexamer cleavage model (see **Methods**). The significance of difference between the observed and expected cleavages is evaluated per nucleotide using the negative binomial dispersion model. Individual *p*-values are combined in 7-bp windows using the Stouffer's Z-score method. Per-nucleotide false discovery rates are computed by sampling from the expected null distributions. **b,** Autocorrelation of *p*-values sampled from the expected negative binomial distribution. **c–d**, Histogram of windowed *p*-values windowed p-values observed (**c**) and sampled (**d**) data. **e,** Observed and sampled *p*-values are compared to empirically determine and calibrate false-positive rates.

##### Extended Data Figure 3. Genomic footprints are reproducible. Harbor evolutionary constrained nucleotides, and overlap TF recognition sequences

**a,** Scatter plot of per-nucleotide footprint *p*-values for replicate experiments from the same cell line (NAMALWA Burkitt's lymphoma cells). All individual nucleotides within FDR 1% footprints in either replicate were considered for correlation analysis. **b,** Same as in **a**, but for replicates of the same primary cell (CD8+ T cells) between two distinct individuals. **c,** Pearson's correlation between replicates pairs grouped by whether they were derived from the same cell and individual ($n$=43) or were the same primary cell or tissue from different individuals ($n$=111). **d,** Evolutionary conservation is associated with footprint confidence. Plotted is per-nucleotide evolutionary conservation (phyloP) (windowed mean) for nucleotides ranked by footprint *p*-value from CD19+ B cells. **e,** Same as in **d**, but for motif overlap.

**Extended Data Figure 4. Empirical Bayes framework identifies footprints with markedly increase reproducibility**

**a,** Histogram of footprint discovery concordance between replicates (same cell type, same individual) using footprints called either independently (FDR 1%) (light orange) or with the Empirical Bayes approach (posterior probability > 0.99).

**Extend Data Figure 5. Clustering motifs by similarity**

**a,** Outline of motif clustering approach. Motif models (n=2,179) from Jolma et al.[9], JASPAR[10] (2018), and HOCOMOCO[11] (version 11) were clustered using pair-wise similarity scores using TOMTOM[14]. **b,** Hierarchically clustered heatmap of the pairwise similarity scores between motifs. The cluster dendrogram was cut at height 0.7 to create non-redundant archetypal clusters of motifs**. c,** Exemplar clusters of similar TF recognition sequences corresponding to KLF/SP (C2H2 family), EGR (C2H2 family), MEF2 (MADS) and E-box/CATATG (bHLH).

**Extended Data Figure 6. Aggregate DNase I cleavage profiles for diverse transcription factors.**

Per-nucleotide DNase I cleavage patterns surrounding instances of motifs within genomic footprints for (**a**) RELA (CD14+ monocytes) (**b**) EBF1 (bipolar neuron), (**c**) PAX6 (fetal eye), (**d**) SOX3 (differentiated neuronal cell), (**e**) MYF6 (fetal tongue) and (**f**) HNF1A (tubular kidney cells). For each panel, top left shows a randomly ordered heatmap map showing the per-nucleotide relative DNase I cleavage protection (observed/expected) for each footprinted motif instance. Below, aggregate DNase I protection averaged over all footprinted sites. Left, DNase I cleavage at individual motif instances (blue, observed cleavage; yellow, expected cleavage).

**Extended Data Figure 7. Cell-selective occupancy of TF recognition sequences**

**a,** Hierarchically clustered heatmap of TF recognition sequence enrichment ($-\log_{10}$ *q*-values) (see **Methods**) overlapping consensus footprints. Rows correspond to motifs and columns correspond to individual samples. **b,** Clustered heatmaps of posterior probabilities for footprints (left) overlapping an E-box/CAGCTG (MYF6_bHLH_1 motif model) and their corresponding DNase I density (right) in each sample. Rows and columns are clustered on footprint probabilities using *K*-means (footprints, *k*=6) and hierarchical (biosamples) clustering. **c–d,** Same as **b**, for footprints overlapping an E-box/CATATG (c, Neurog1_MA0623.1 motif model) or MEIS (d, MEIS1_MEIS_1 motif model) recognition sequence.

**Extended Data Figure 8. Differential occupancy within the UCP2 promoter in human brain and neuronal cells**

Comparative footprinting within the *UCP2* promoter identifies sequences differentially occupied in neuronal and non-neuronal cell and tissue types. Top, DNase I cleavage in two neuronal and non-neuronal cell types. Bottom, differential footprint test between 28 nervous- and 189 non-nervous related biosamples identifies differentially footprinted nucleotides.

#### Extend Data Figure 9. Differentially occupied nucleotides reflect aggregate DNase I cleavage profiles

**a,** Density histograms of relative footprint occupancy between nervous-system derived and non-nervous-system derived samples for the TF recognition sequences of REST, NFIB, ZIC1 and EBF1. Grey indicates distribution of all motif instances tested. Black indicates differentially footprinted. **b,** Per-nucleotide aggregate plots of the mean relative DNase I protection (top) and differential test p-value ($-\log_{10}$, bottom) around differential occupied motifs. **c,** Aggregate DNase I cleavage around all footprints containing the same recognition sequences as **a** from either nervous- or non-nervous-system related cells/tissues. **d,** Absolute Pearson's correlation coefficient of the aggregated relative protection (**b**, top) and DNase I cleavage profiles (**c**) for each TF.

#### Extended Data Figure 10. Expression of genes nearby differentially occupied REST sites is enriched in brain and neuronal cell types

For each differentially occupied REST site, the two closest genes were identified and cell and tissues enrichment was performed using Enrichr.[26] Shown are significantly enriched cell and tissues (adjusted p-value<0.01).

#### Extended Data Figure 11. Detection and quantification of allelic imbalance

**a,** Scatterplot of allelic ratios at 100 randomly selected high confidence SNVs (see **Methods**) computed after aggregating reads from different samples (*x*-axis) vs. the distribution of allelic ratios at the same SNVs in each sample (*y*-axis; mean ± s.d.). The average standard deviation is indicated in the top left corner as is used to tune the parameters of a Beta-binomial distribution. **b,** Simulation of allelic ratios rom the observed total read depth at high confidence SNVs assuming a binomial distribution ($p$=0.5) or a Beta-binomial distribution. Grey indicates the observed allelic ratios at the same variants. **c,** Density histogram of allelic ratios for all tested SNVs (grey line) and significantly imbalanced SNVs (blue line).

#### Extended Data Figure 12. Allelically imbalanced variants are enriched within consensus footprints.

SNVs within consensus footprints (posterior probability > 0.99) have increased frequency of imbalance irrespective of read depth.

**Extended Data Figure 13. Enrichment of imbalanced variants within footprinted TF recognition sequences.**

**a–d,** Distribution of SNVs around the recognition sequences for CREB/ATF, NFI, TEAD and TFAP2 transcription factors. For each TF, shown are the total SNVs tested for imbalance (top), imbalanced variants (middle), and the proportion of variants imbalanced stratified by whether they additionally overlap a consensus footprint. **e,** $\log_2$ enrichment of imbalanced variants residing within TF recognition sequences relative to non-imbalanced SNVs for both variants within footprinted (blue) and non-footprinted motifs (orange). Motifs are grouped into clusters, where each point represents an individual motif model (see **Extended Data Fig. 5**, **Extended Data Table 2** and **Methods**). Black bars indicate the mean enrichment across all motifs in each cluster and footprint overlap. Only motifs with significant (q-value<0.05) enrichment of imbalanced SNVs with a footprinted recognition sequence are shown.

**Extended Data Figure 14. Allelic imbalance parallels the predicted energetic effect of the variant on the TF binding site**

**a,** Density histogram of allelic ratios for variants overlapping footprinted CREB1 (CREB/ATF), TEAD1 (TEAD) and TFAP2C (TFAP2) recognition sequence. Grey line, all variants tested for imbalance. Blue line, all variants significantly imbalanced. **b,** Shown is the mean log-odds motif score (reference vs. alternate allele) of all tested variants within footprinted motifs binned by allelic ratios.

**Extended Data Figure 15. Disease- and trait-associated variation is enriched within genomic footprints**

Observed and expected colocalization of EBI/NHGRI GWAS Catalog SNVs to DNase I hypersensitive sites and consensus footprints. Grey histogram shows the number of overlapping SNPs from 1,000 sets of matched sampled variants from the 1000 Genomes Project (CEU population), with the black dashed line indicating the mean. Red arrow indicates the number of overlapping GWAS variants.

#### Extended Data Tables

**Extended Data Table 1. Overview and summary of DGF data used in this study**

**Extended Data Table 2. Clustering of TF recognition sequence models**

#### Extended Data Files

**Extended Data File 1. Footprints identified in individual datasets**

BED-formatted files contain coordinates of footprints called in individual biosamples (FDR<0.01)

**Extended Data File 2. Consensus-defined footprints across 243 biosamples**

BED-formatted file containing coordinates of consensus footprints (GRChr38 reference genome) called at a posterior probability > 0.99.

Column definitions:

1. Chromosome
2. Start (0-based)
3. End
4. Unique identifier
5. Motif cluster matches (format: [cluster number]:[seed motif], semi-colon delimited) (see *Extended Data Table 2* for cluster information)

**Extended Data File 3. Posterior probability matrix of consensus footprints by biosamples**

Matrix of posterior probabilities across all biosamples (columns) and consensus footprints (rows). Rows correspond to footprints (same order and identifiers as *Extended Data File 2*, columns correspond to biosamples (same order as *Extended Data Table 1*).

#### Extended References


1. Thurman, R. E. *et al.* The accessible chromatin landscape of the human genome. *Nature* **489**, 75–82 (2012).

2. Consortium, R. *et al.* Integrative analysis of 111 reference human epigenomes. *Nature* **518**, 317–330 (2015).

3. John, S. *et al.* Genome-scale mapping of DNase I hypersensitivity. *Current protocols in molecular biology* **Chapter 27**, Unit 21.27-21.27.20 (2013).

4. Li, H. & Durbin, R. Fast and accurate short read alignment with Burrows-Wheeler transform. *Bioinformatics (Oxford, England)* **25**, 1754–1760 (2009).

5. Vierstra, J. & Stamatoyannopoulos, J. A. Genomic footprinting. *Nature methods* **13**, 213–221 (2016).

6. Lazarovici, A. *et al.* Probing DNA shape and methylation state on a genomic scale with DNase I. *Proc. Natl. Acad. Sci. U. S. A.* **110**, 6376–6381 (2013).

7. Meuleman, W. *et al.* Index and biological spectrum of accessible DNA elements in the human genome. *Biorxiv* 822510 (2019) doi:10.1101/822510 .

8. Cooper, G. M. *et al.* Distribution and intensity of constraint in mammalian genomic sequence. *Genome research* **15**, 901–913 (2005).

9. Jolma, A. *et al.* DNA-binding specificities of human transcription factors. *Cell* **152**, 327–339 (2013).

10. Khan, A. *et al.* JASPAR 2018: update of the open-access database of transcription factor binding profiles and its web framework. *Nucleic acids research* **46**, D1284 (2018).

11. Kulakovskiy, I. V. *et al.* HOCOMOCO: towards a complete collection of transcription factor binding models for human and mouse via large-scale ChIP-Seq analysis. *Nucleic acids research* **46**, D252–D259 (2018).

12. Korhonen, J. H., Palin, K., Taipale, J. & Ukkonen, E. Fast motif matching revisited: high-order PWMs, SNPs and indels. *Bioinformatics (Oxford, England)* **33**, 514–521 (2017).

13. Maurano, M. T. *et al.* Large-scale identification of sequence variants influencing human transcription factor occupancy in vivo. *Nature genetics* **47**, 1393–1401 (2015).

14. Gupta, S., Stamatoyannopoulos, J. A., Bailey, T. L. & Noble, W. Quantifying similarity between motifs. *Genome biology* **8**, R24 (2007).

15. Lei, X. *et al.* The Cancer Mutation D83V Induces an α-Helix to β-Strand Conformation Switch in MEF2B. *Journal of molecular biology* **430**, 1157–1172 (2018).

16. Yin, M. *et al.* Molecular mechanism of directional CTCF recognition of a diverse range of genomic sites. *Cell research* **27**, 1365–1377 (2017).

17. Schr\"odinger, L. The PyMOL Molecular Graphics System, Version~1.8. (2015).

18. Geijn, B., Mcvicker, G., Gilad, Y. & Pritchard, J. K. WASP: allele-specific software for robust discovery of molecular quantitative trait loci. bioRxiv. (2014).

19. Buniello, A. *et al.* The NHGRI-EBI GWAS Catalog of published genome-wide association studies, targeted arrays and summary statistics 2019. *Nucleic acids research* **47**, D1005–D1012 (2019).

20. Bulik-Sullivan, B. K. *et al.* LD Score regression distinguishes confounding from polygenicity in genome-wide association studies. *Nature genetics* **47**, 291–295 (2015).

21. Finucane, H. K. *et al.* Partitioning heritability by functional annotation using genome-wide association summary statistics. *Nature genetics* **47**, 1228–1235 (2015).

22. Neph, S. *et al.* BEDOPS: high-performance genomic feature operations. *Bioinformatics (Oxford, England)* **28**, 1919–1920 (2012).

23. Zhao, H. *et al.* CrossMap: a versatile tool for coordinate conversion between genome assemblies. *Bioinformatics (Oxford, England)* **30**, 1006–1007 (2014).

### Extended Data Figure 1

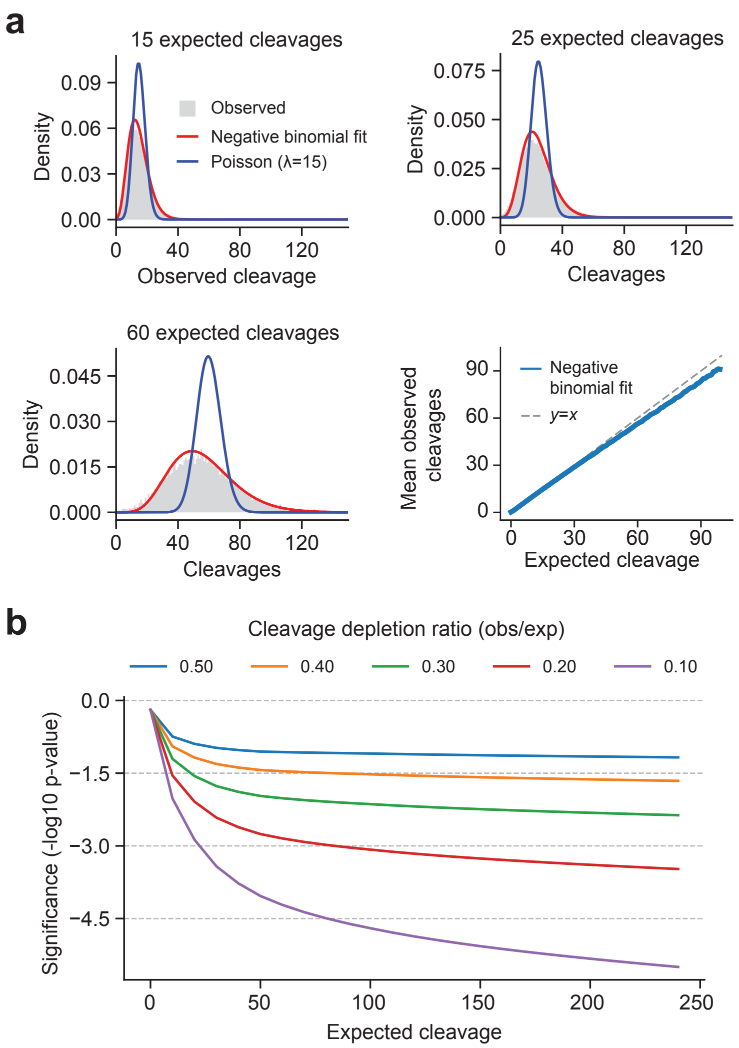

### Extended Data Figure 2

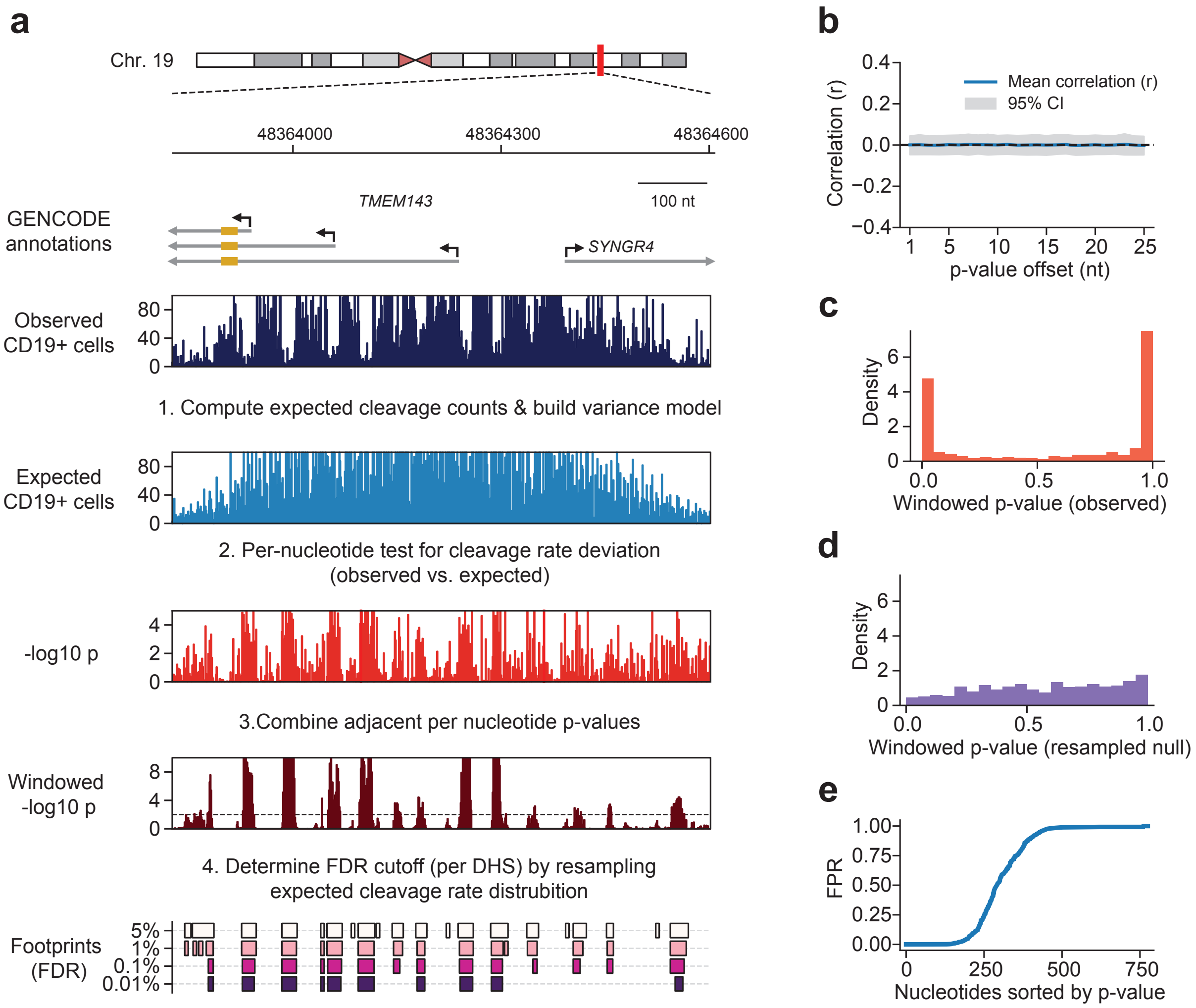

### Extended Data Figure 3

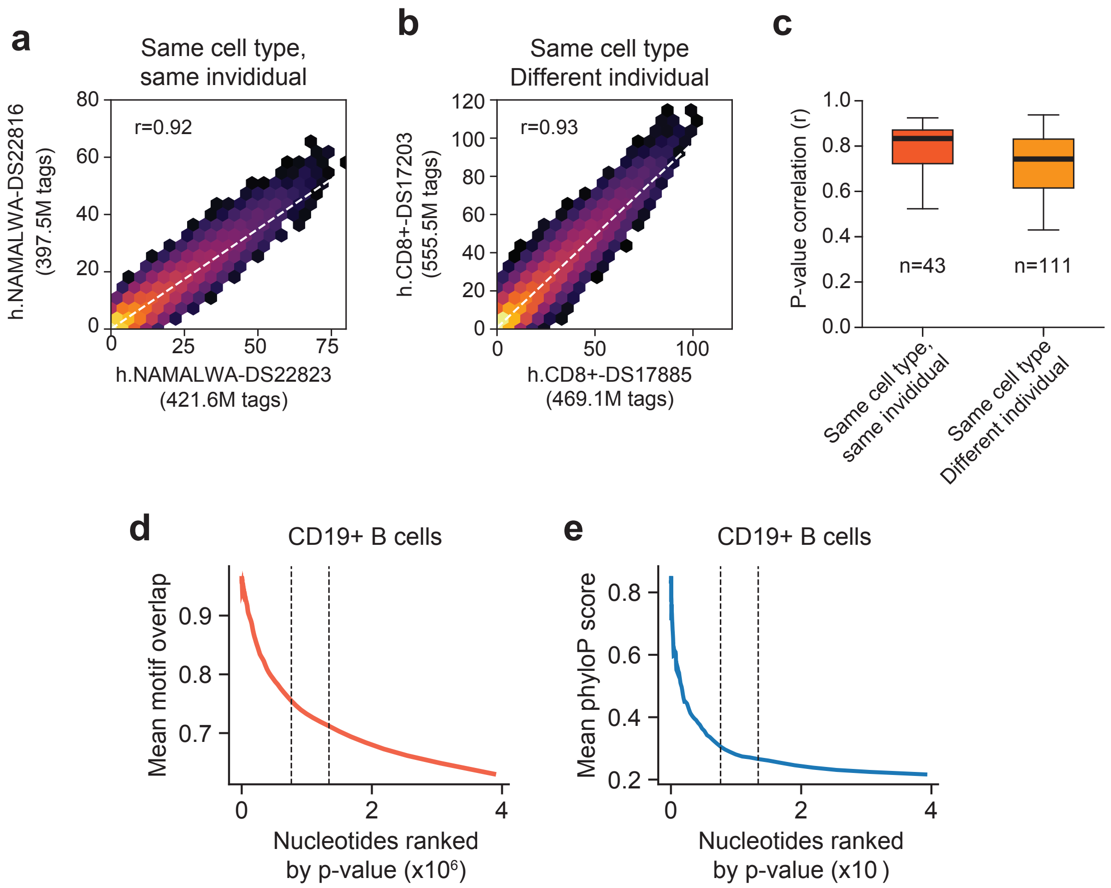

### Extended Data Figure 4

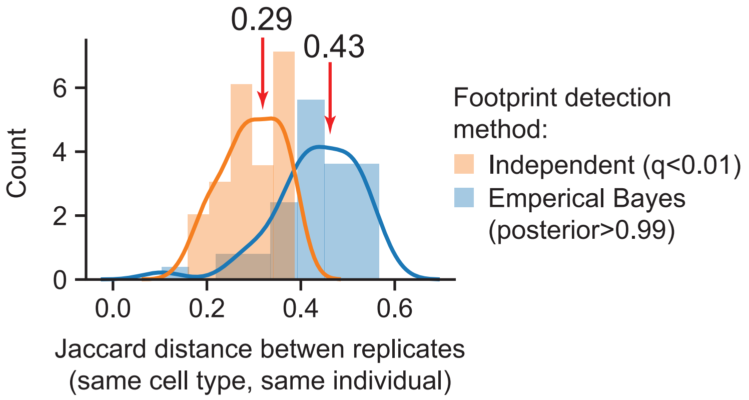

### Extended Data Figure 5

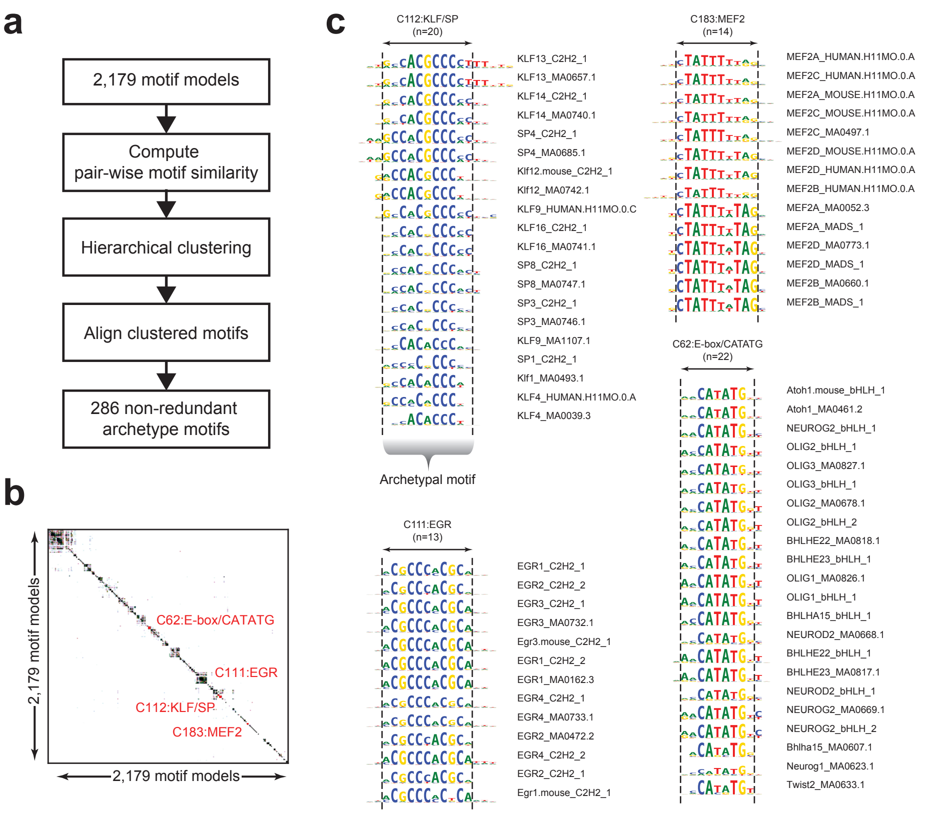

### Extended Data Figure 6

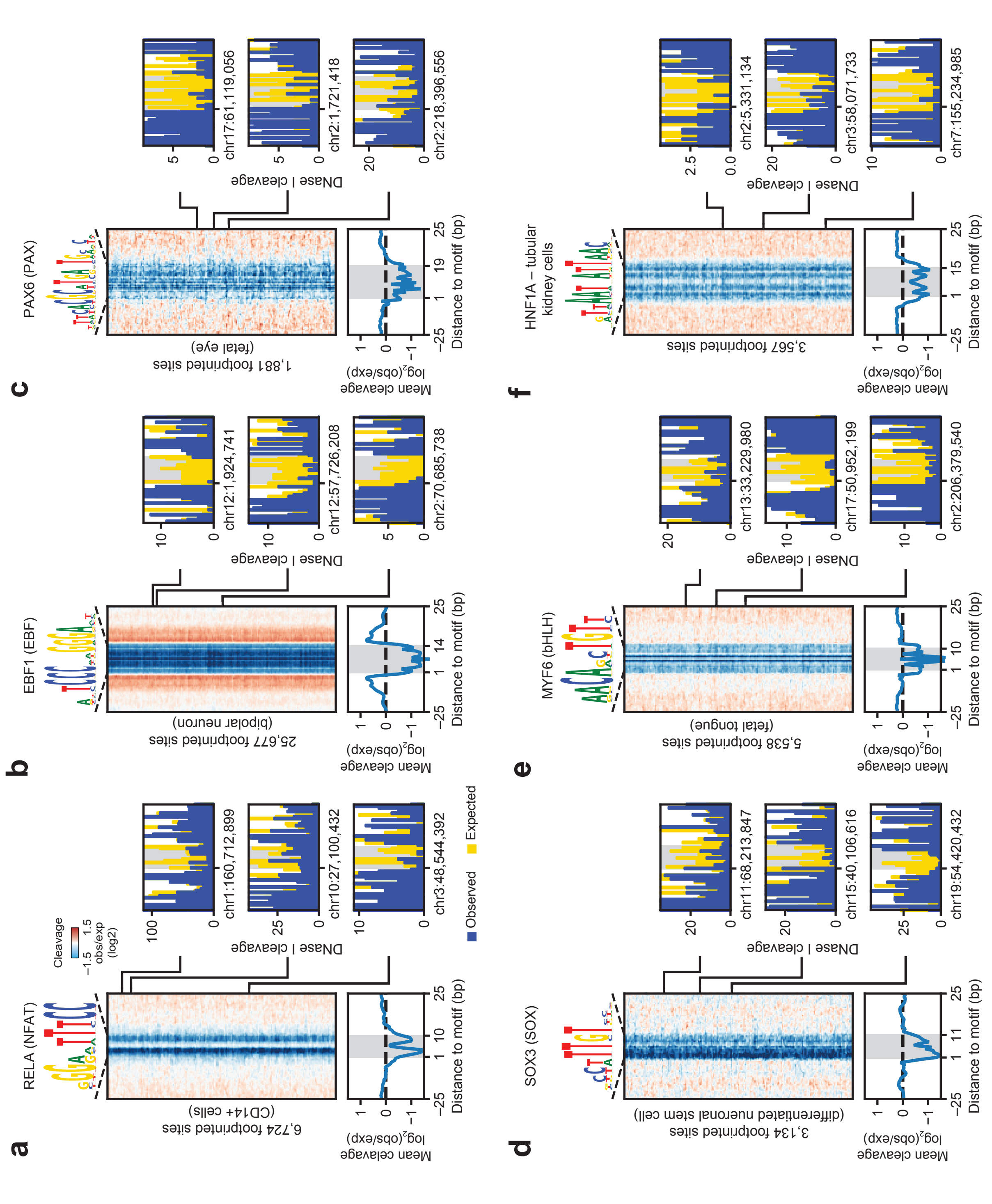

### Extended Data Figure 7

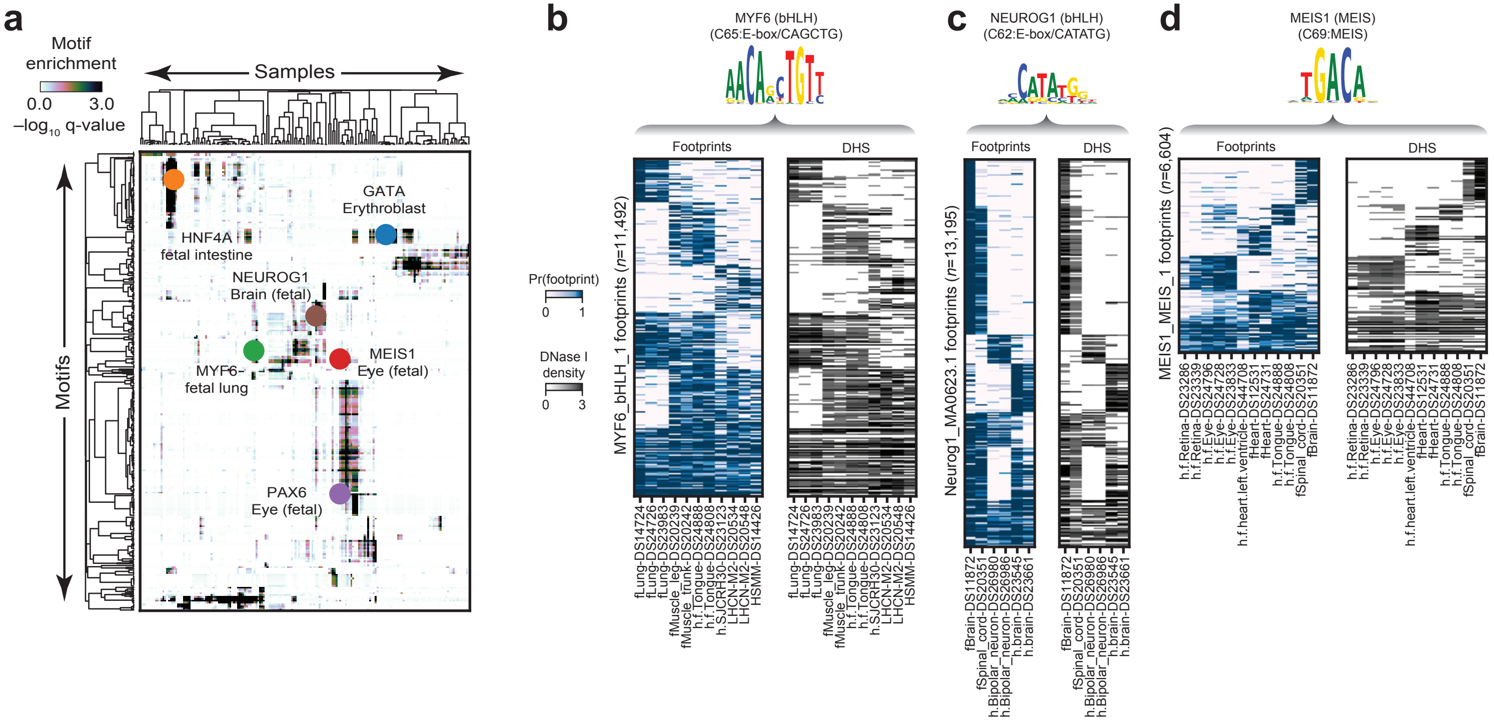

### Extended Data Figure 8

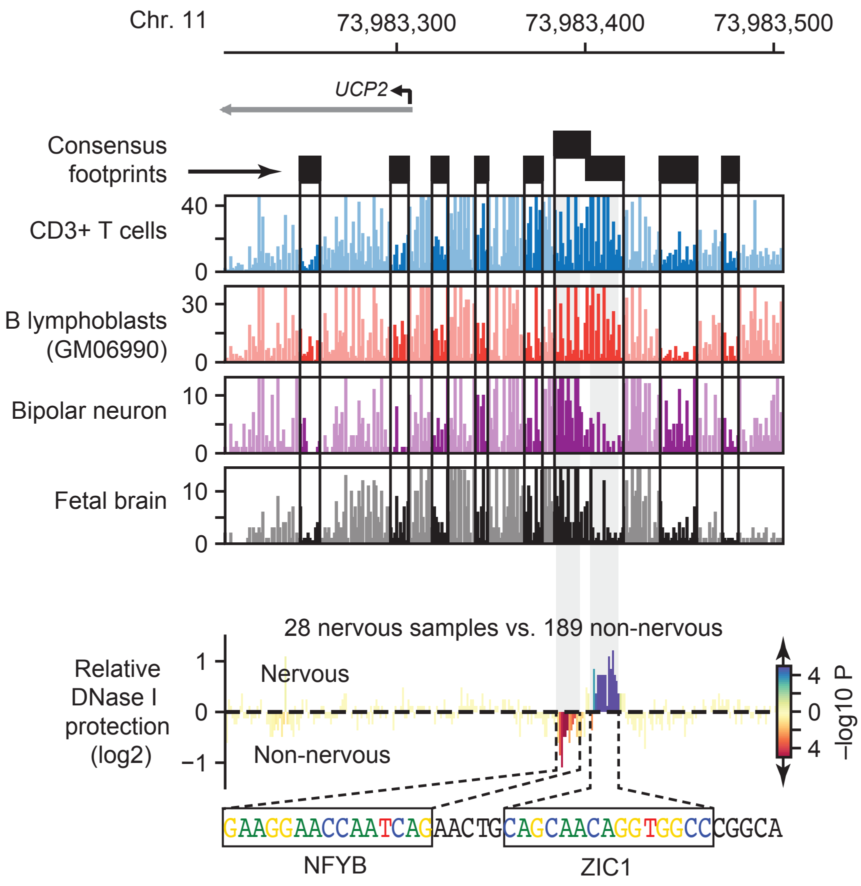

### Extended Data Figure 9

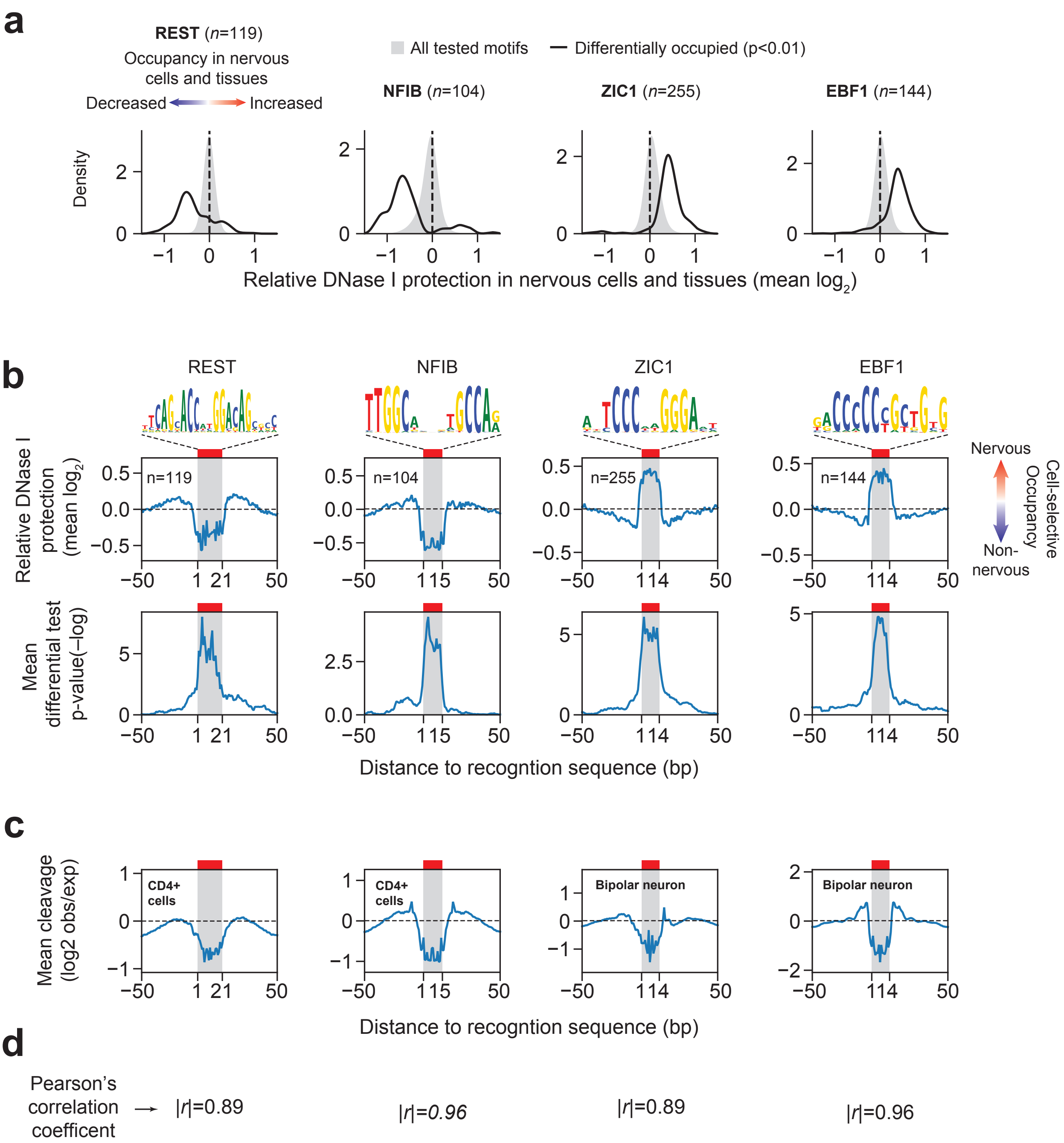

### Extended Data Figure 10

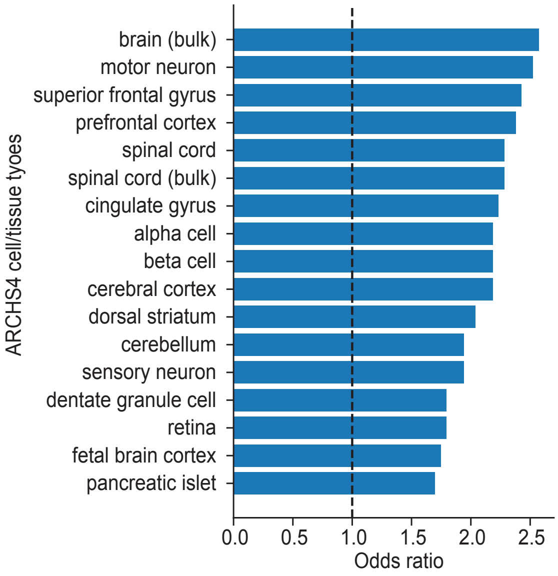

### Extended Data Figure 11

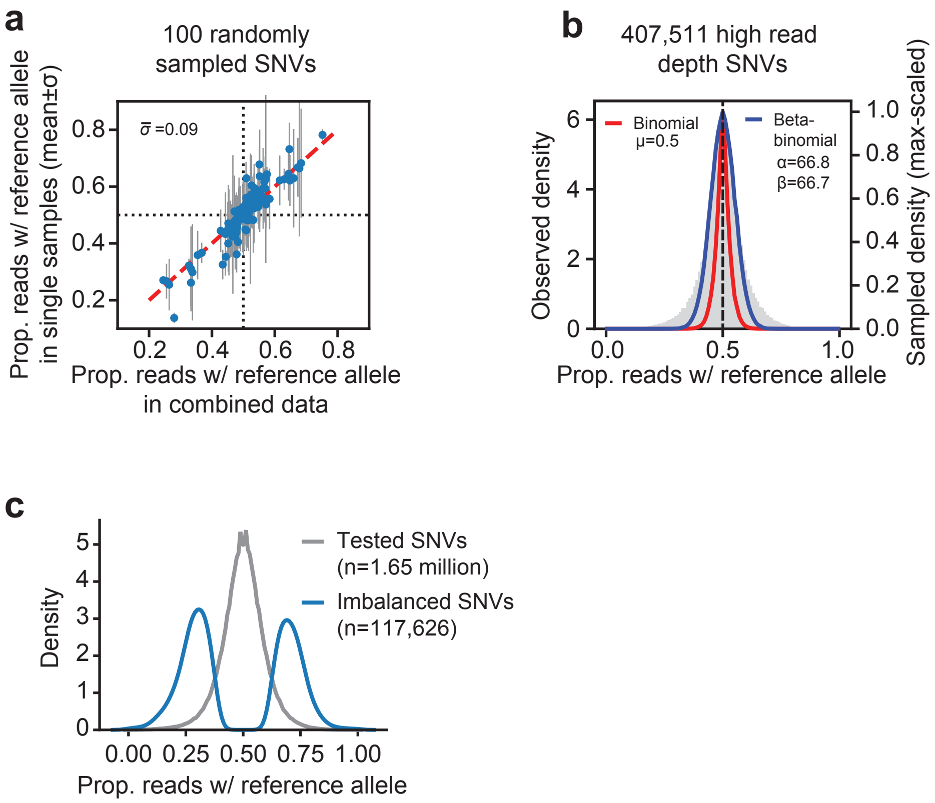

### Extended Data Figure 12

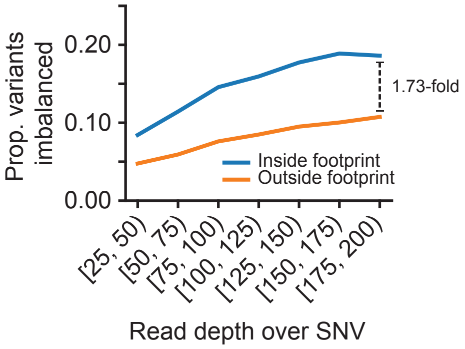

### Extended Data Figure 13

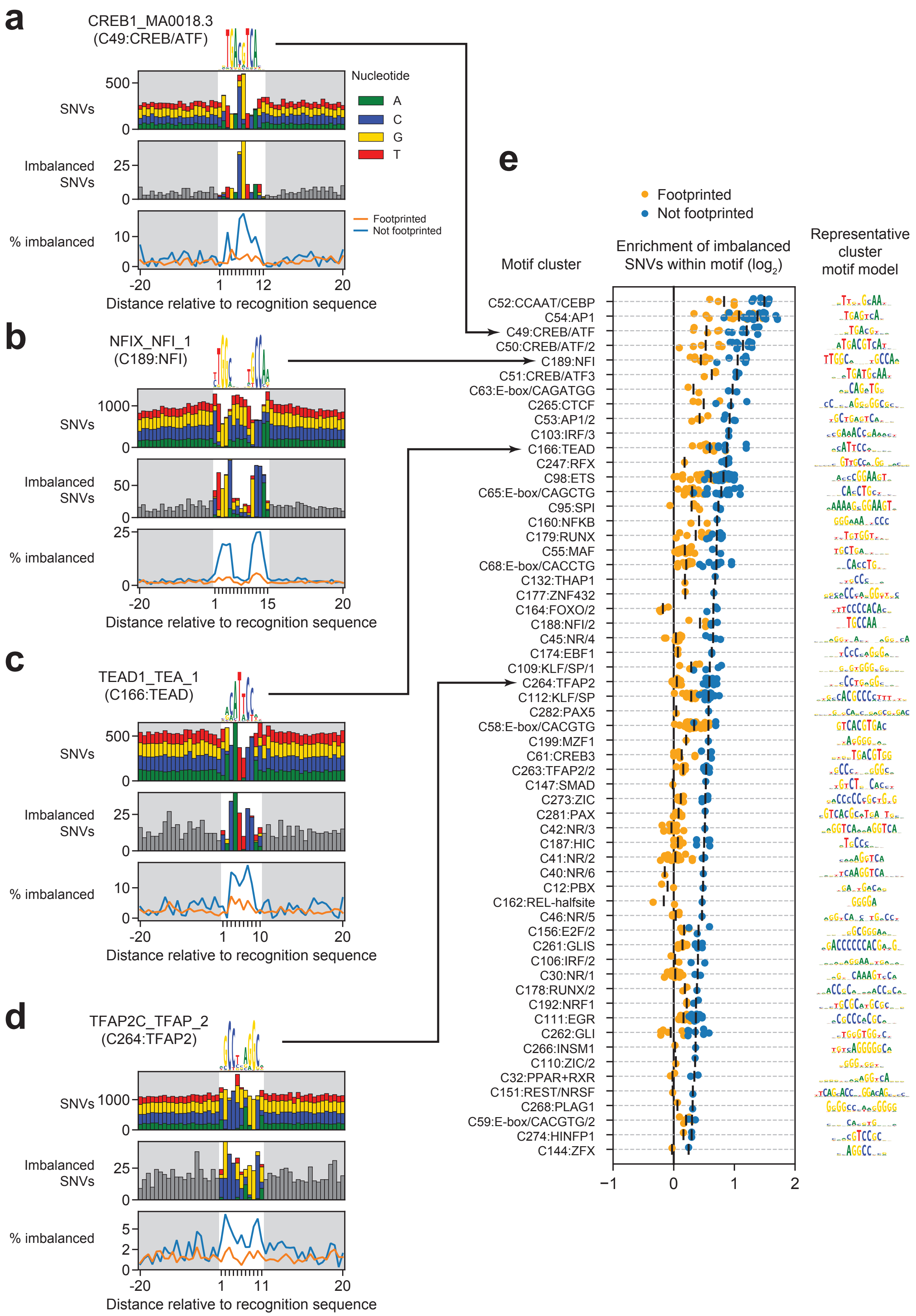

### Extended Data Figure 14

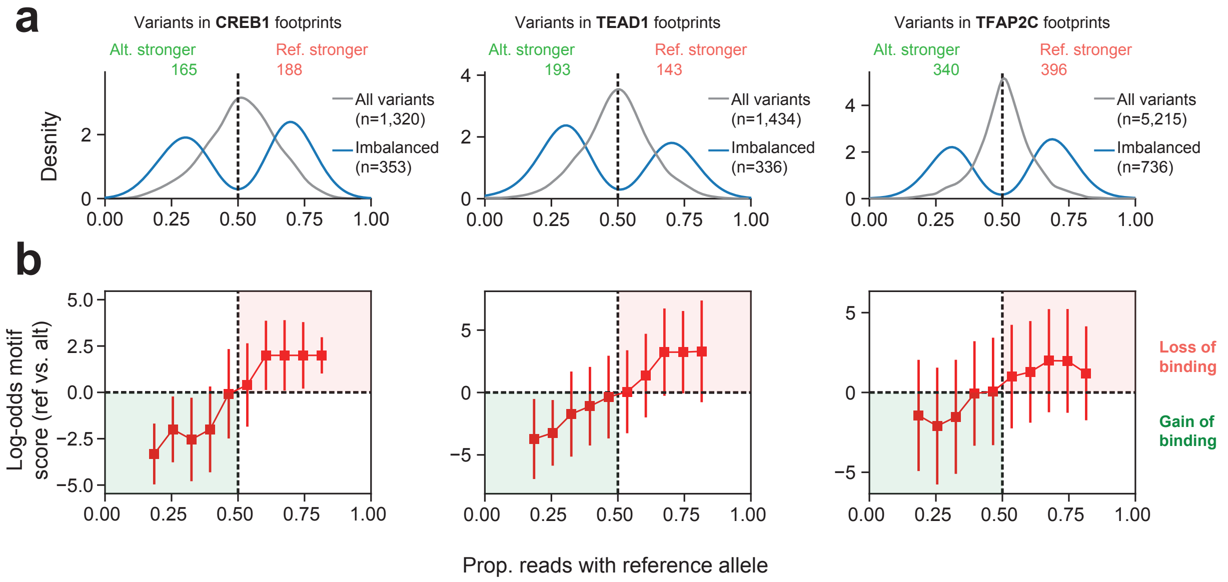

### Extended Data Figure 15

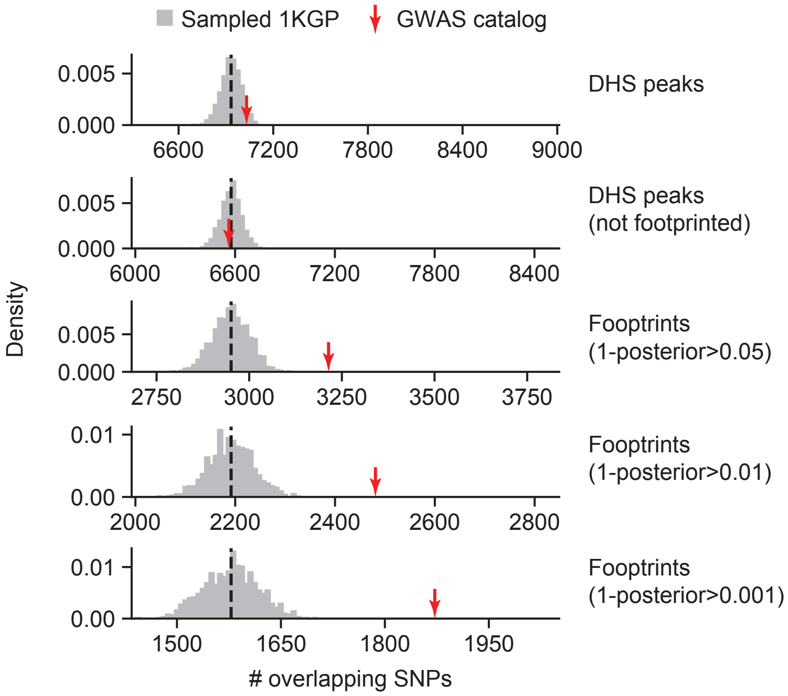
